# Wild and farmed *Saccharina latissima* in Europe: genetic insights for sustainable cultivation, traceability and environmental challenges

**DOI:** 10.1101/2025.02.06.636791

**Authors:** Lucie Jaugeon, Christophe Destombe, Paolo Ruggeri, Stéphane Mauger, Jérôme Coudret, J. Mark Cock, Philippe Potin, Myriam Valero

## Abstract

Kelp cultivation is expanding rapidly worldwide, although the aquaculture of *Saccharina latissima* (sugar kelp) in Europe remains in its early stages. However, major concerns have emerged about the potential impact of selected cultivars of the species on native populations of *S. latissima* which are already vulnerable to stressors including climate change.

To address these concerns and support sustainable cultivation, it is essential to characterise genetic diversity and structure of wild kelp populations to monitor potential farm-to-wild gene flow. In this study we used 21 microsatellite loci to characterise the genetic structure of 24 natural and 3 farmed populations of *S. latissima* along European coasts. Results confirmed strong genetic differentiation between the northern and southern coasts, refining the boundaries between these two clusters compared to previous studies. Within each cluster, a clear genetic substructure was detected, with population differentiation being far more pronounced in the southern cluster.

Cultivated sporophytes exhibited pedigrees that were all traced back to the local parent populations. Bayesian model-based structure analysis, discriminant analysis of principal components and assignment tests revealed no significant genetic differentiation between the farms and their wild populations. This indicates that farmed populations and neighbouring populations share the same gene pool, reflecting current cultivation practices. These findings contribute to understanding risks of gene flow between wild and farmed populations and demonstrate that strain traceability is feasible. Additionally, the study highlights the challenge European seaweed farmers face in securing reliable seedling stock, especially in the context of global change.

## INTRODUCTION

The increasing diversification of seaweed applications in industries such as food, pharmaceuticals, agriculture, textiles, biodegradable materials, is driving a major transition in the way these resources are exploited. To ensure sustainability, the harvesting of wild populations is now being supplemented or replaced by intensive cultivation (Buschmann et al., 2014; Brakel et al., 2021). Over the past 22 years, the global production of cultivated marine macroalgae - or seaweeds - has increased from 12 to 36 million of wet tons, now representing 51% of total marine aquaculture production, while harvesting from wild populations remains at just 1 million wet tons (FAO, 2024). New aquaculture techniques are emerging to meet the growing demand (Hafting et al., 2015; Garcia-Poza et al., 2020), and kelp farming especially faces a considerable challenge to increase production around the world (Ferdouse et al., 2018; FAO, 2020).

However, the expansion of intensive open-sea cultivation can have significant and sometimes irreversible consequences on the environment. In particular, the dispersal of cultivated genotypes and possible hybridization of farmed seaweeds with native stocks can result in impoverishment of local genetic diversity (Hutchings & Fraser, 2008; Liu et al., 2012). The major risk associated with aquaculture is the introduction of cultivated organisms into wild populations. Furthermore, the transfer of stocks around the world increases the risk of diseases and pests (Naylor et al., 2001; Naylor et al., 2021). The introgression of farmed genetic material into wild populations has been extensively documented in terrestrial agriculture and animal aquaculture (see for review: Manchester & Bullock, 2000; Ellstrand et al., 2013); however, it remains poorly understood in the context of seaweed aquaculture (Valero et al., 2017; Goecke et al., 2020). This “genetic pollution” could profoundly affect the genetic diversity and evolutionary trajectory of seaweed natural populations (Loureiro et al., 2015, Valero et al., 2017). Indeed, in species where natural populations are highly structured geographically, care should be taken to not translocate individuals among genetically isolated populations (Nepper-Davidsen, 2021). Additionally, cultivated individuals may benefit from a selective/demographic advantage compared to individuals from native populations and spread in the natural environment, reducing biodiversity (Williams and Smith, 2007).

The sugar kelp *Saccharina latissima* (Linnaeus) C.E. Lane, C. Mayes, Druehl & G.W. Saunders is a promising candidate for intensive aquaculture due to its rapid growth and diverse industrial applications (Sæther et al., 2024). In order to get improved strains for the *S. latissima* cultivation industry, different breeding programs have been initiated and in particular the identification of temperature-stress-related quantitative trait loci (Huang et al., 2023; Goecke et al., 2020; Nehr et al., in prep.). This species, which spans the northern hemisphere from polar to temperate regions, is one of the most extensively studied kelps in terms of its physiology and ecology (see for review: Bartsch et al., 2008; Diehl et al., 2024). Generally, dispersal in kelp is typically limited, with recruitment occurring within a few metres of parental sporophytes (Santelices, 1990; Kinlan & Gaines, 2003). Molecular studies (COI and microsatellite markers) have revealed that *S. latissima* represents a complex of species, comprising the north-east Pacific *S. cichorioides* and distinct lineages in the North Atlantic (Neiva et al., 2018). In Europe, based on these markers and SNPs, a single phylogroup stretches from Spitsbergen to Portugal, showing clear genetic differentiation between northern and southern Europe (Neiva et al., 2018; Guzinski et al., 2020). While significant genetic structure has been observed between distant populations (Neiva et al., 2018; Guzinski et al., 2016, 2020; Evankow et al., 2019; Ribeiro et al., 2022), *S. latissima* generally shows lower genetic diversity and differentiation compared to other European kelps like *Laminaria digitata* (Billot et al., 2003; Neiva et al., 2020) and *L. hyperborea* (Robuchon et al., 2014; Evankow et al., 2019). This suggests that dispersal in *S. latissima* may be more significant than in other kelp species, raising questions about the potential extent of gene flow between cultivated and native populations. Interestingly, in Asia, *S. japonica* farms showed minimal gene flow to adjacent wild populations, likely due to cultivation practices such as selective breeding and pre-fertility harvesting (Shan et al., 2019).

So far, small-scale *S. latissima* cultivation trials have been conducted across Europe, from the Iberian Peninsula (Peteiro & Freire, 2013; Freitas et al., 2015) to Norway (Handa et al., 2013; Forbord et al., 2020), as well as in Brittany (Mesnildrey et al., 2012), Ireland (Mac Monagail & Morisson, 2020), Scotland (Capuzzo & McKie, 2016), the Netherlands (Van Oirschot et al., 2017), Denmark (Boderskov et al., 2021; 2023), Faroe Islands (Bak, 2019) and Sweden (Visch et al., 2020). These small farms typically seed their crop lines using a mixture of sporophytes broodstock from local indigenous populations (Araujo et al., 2021). Open-water cultivation usually takes a few months with cultivated sporophytes being harvested before reaching fertility. However, gene flow from farmed to wild populations remains a potential risk if spores are released prematurely or if sporophytes remain on the lines after harvesting or detach from the lines.

In order to assess the origin of seedlings and the level of connectivity between farms and natural populations in Europe, it is necessary to have a good knowledge of the genetic landscape of the study species in this region. The first objective of this study was to refine the pattern of genetic structure along the European coast, particularly focusing on the boundaries between the northern and southern European groups of *S. latissima* using 21 microsatellite markers across 24 natural populations. The second objective was to evaluate the genetic diversity of three cultivated populations and compare them to local wild populations to evaluate strain traceability and potential gene flow risks associated with farming practices.

## MATERIALS AND METHODS

### Sample collection

Sporophytes of *S. latissima* were collected from 27 localities (24 natural populations, N = 677 and 3 farms, N = 101) across the North-Eastern Atlantic, between 2017 and 2019 (Figure 1 and Table 1). For each sampling campaign, the national regulations pertaining to the Nagoya Protocol were followed. For the farms, sporophytic individuals were randomly harvested from aquaculture lines. In two of the three farms, the exact source population used to seed the farms was also sampled: natural population 9 (Saint-Brieuc) for farm 25 (C-WEED) and natural population 24 (Frøya) for farm 27 (SES). For farm 26 (SAMS), the exact natural population used to seed this farm is known but could not be sampled, instead we sampled natural population 16 (Atlantic Bridge) for comparison with the farm. Population 16 is approximately 20 km from the true natural population used to seed the SAMS farm. In natural populations, sampling was carried out on individuals attached to the substrata in the lower intertidal zone except for the two localities of Piriac-sur-Mer and Noirmoutier (Figure 1) where drifting individuals were sampled along the beach. For both farms and natural populations, a small disc of tissue (2–4 cm^2^) was excised from the base of each individual’s blade. The dried tissue samples were stored in individual zip-locked plastic bags with silica gel for preservation until DNA extraction.

**Fig. 1.**
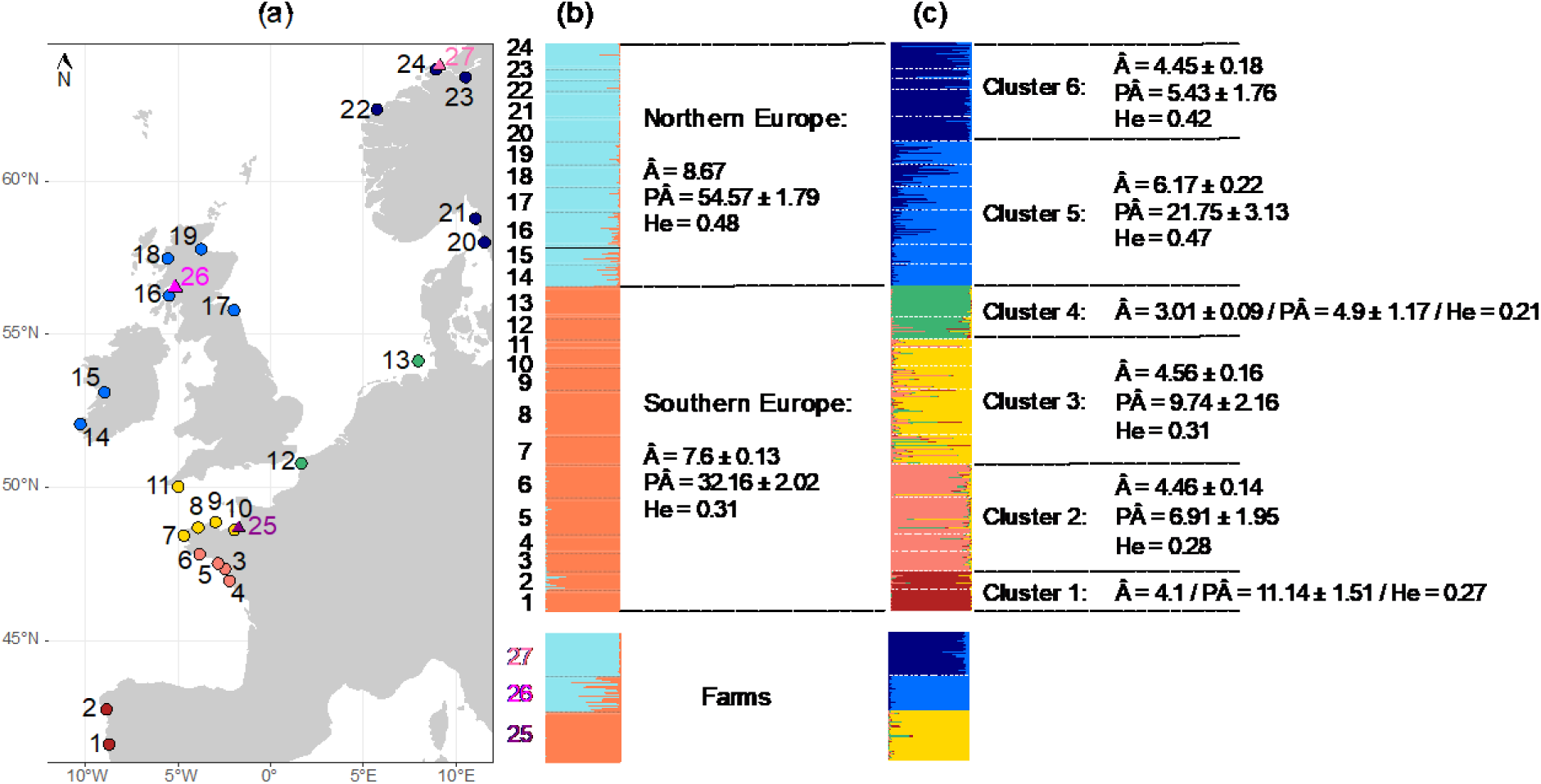
Sampling locations and genetic structure of *Saccharina latissima* based on multilocus microsatellite genotypes. (a) Map of sampling sites, with locations colored according to genetic structure inferred by STRUCTURE. Circles represent natural populations, while triangles indicate farmed populations. (b) Uppermost level of genetic structure, standardised allelic richness (Â), standardised number of private alleles (PÂ) and gene diversity (He) per cluster for a common sample of 256 individuals. Values are represented as means ± standard deviation. (c) Second hierarchical level of structure, Â, PÂ and He per cluster for a common sample of 42 individuals

**Table 1.**
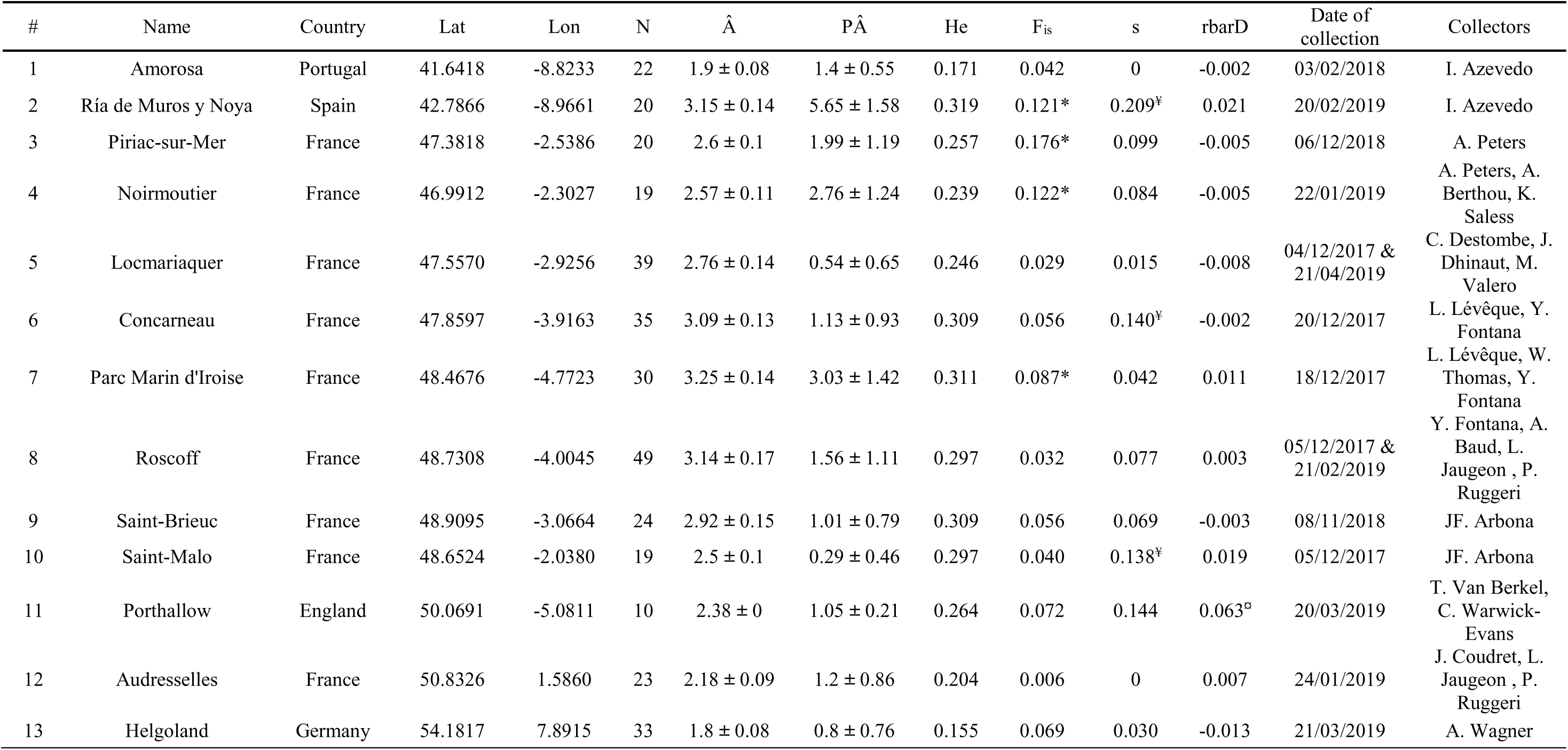
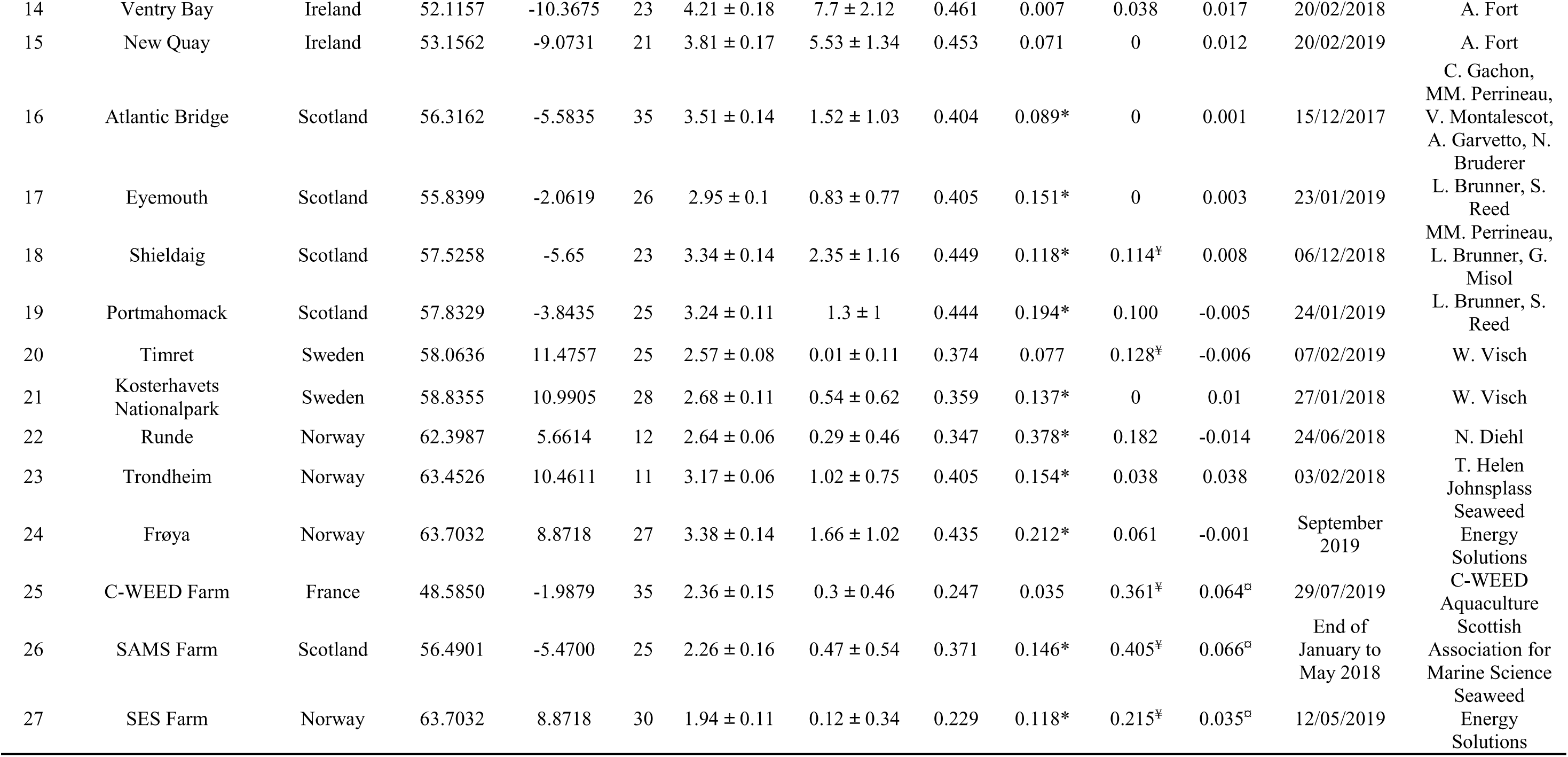
List of samples of *Saccharina latissima*. Site number (#), site name, country, latitude (Lat) and longitude (Lon) in decimal degrees, sample size (N, number of individuals), standardised allelic richness (Â) and standardised number of private alleles (PÂ) for a common sample size of 10 individual (mean ± standard deviation), expected heterozygosity (He), multilocus FIS estimates (* indicates significant deviation from Hardy– Weinberg equilibrium), selfing rate (s) (¥ indicates significant heterozygosity disequilibrium) and modified index of association (rbarD) (¤ indicates significant linkage disequilibrium)

### Microsatellite genotyping

Total genomic DNA was extracted from approximately 10-20 mg of silica gel-dried tissue. DNA extraction was performed using the Nucleospin® 96 plant kit (Macherey-Nagel, Düren, Germany) according to the manufacturer’s instructions, except that cell lysis was performed at 65°C for 15 minutes and the washing steps with buffer PW1 and PW2 were each repeated twice. These modifications were performed to remove most of the polysaccharides that might inhibit DNA extraction and PCR amplification of microsatellite markers (SSRs).

Amplification and scoring of the SSR loci were performed as detailed in Guzinski et al. (2016) for 19 expressed sequence tag (EST)-derived SSR loci, and as in Paulino et al. (2016) for 8 genomic SSR loci. Amplification products were separated by electrophoresis on an ABI 3130 XL capillary sequencer (Applied Biosystems, USA). Alleles were sized using the SM594 size standard (Mauger et al., 2012) and scored manually using the software GeneMapper 4.0 (Applied Biosystems, Foster City, USA). Two filters were applied to the raw multilocus genotype data-set using the poppr R package version 2.9.6 (Kamvar et al., 2014). First, only loci and individuals with less than 30 % of missing data were retained (4 loci and 89 individuals were removed) and, second, loci with global minor allele frequency (MAF) of less than 0.01 were discarded (2 additional loci were excluded). The final dataset included 599 wild and 90 farmed sporophytes characterised with 21 microsatellite loci (all 18 loci used in Guzinski et al., 2020 except Sacl-32 and Sacl-75, and 5 additional markers: Sacl-11/37/54/65/95 from Guzinski et al., 2016; not used in Guzinski et al., 2020).

### Data analysis

Prior to genetic diversity and population structure analysis, loci were tested for stuttering, large allele dropout and null alleles using the software MicroChecker v2.2.3 (Van Oosterhout et al., 2004). The frequency of null alleles per locus and per population was estimated according to the EM algorithm (Dempster et al., 1977) using FreeNA software (Chapuis and Estoup, 2007; https://www1.montpellier.inra.fr/CBGP/software/FreeNA/).

#### Genetic diversity of natural and farmed populations

Genetic diversity was estimated per population as Nei’s gene diversity (expected heterozygosity, He), allelic richness (Â) and number of private alleles (PÂ). These were standardised for the smallest sample sizes in terms of individuals within sites (N = 10), within regions (N = 42) and within clusters (N = 256) defined thereafter, using 1000 randomizations (Assis et al., 2016).

Estimation of fixation index (F_IS_) per population was computed using Genetix 4.05 (Belkhir et al., 2004) and alleles were randomised 10000 times among individuals within each sample to test for deficits in heterozygotes due to potential non-random mating. The threshold for significant tests was set at p-value < 0.01.

Heterozygote deficiencies often occur in natural populations even when there is limited selfing, due to technical artefacts such as null alleles (Pompanon et al., 2005). To account for technical biases, population selfing rate “s”, derived from “g2”, was calculated using the RMES software (robust multilocus estimate of selfing) (David et al., 2007; https://www.cefe.cnrs.fr/images/stories/DPTEEvolution/Genetique/fichiers%20Equipe/RMES%202009%282%29.zip). This index g2 measures the extent to which heterozygosities are correlated across loci. Under no inbreeding, statistical independence of the heterozygosities at different loci is expected. To test the null hypothesis of no heterozygosity disequilibrium (g2=s=0), 1000 permutations of the genetic data were performed.

Linkage among loci is expected to be very low or null at all SSRs when populations are sexually reproducing because alleles recombine freely to form new genotypes during the process of sexual reproduction. To test this phenomenon, linkage disequilibrium (LD) was assessed per population, using a single multilocus measurement of LD, rbarD, a modified index of association which accounts for unequal sample size (Brown et al., 1980 modified by Agapow and Burt, 2001). rbarD was computed using the poppr R package v2.9.6 (Kamvar et al., 2015). 1000 permutations of the genetic data were used to test the null hypothesis of no linkage among the different loci.

#### Genetic structure of natural populations and assignment of farmed sporophytes

The potential effect of null alleles on genetic differentiation was checked by estimating pairwise *F*_st_ of Weir (1996) both using and without using the ENA correction implemented in the software FreeNA and described in Chapuis and Estoup (2007). The uncorrected and corrected pairwise *F*_st_ were then compared using a paired t-test.

The dataset was used to perform a series of statistical analyses that aimed to detect the number of genetic clusters composed by the different populations of *S. latissima* sampled in our study. The information concerning the genetic structure of *S. latissima* across Europe was then used to perform an assignment test to detect unexpected gene flow between populations and between wild populations and farms.

Genetic structure was inferred using STRUCTURE version 2.3.4 (Pritchard et al., 2000) with admixture and a correlated allele frequency model, without any prior population assignments. STRUCTURE was run in a hierarchical way, evaluating first all populations, and then within the two main clusters South and North Europe separately. A range of assumed populations (K, set sequentially from 1 to 16 for all populations, from 1 to 13 for South Europe, and from 1 to 11 for North Europe) was run 10 times using a burn-in of 5 × 10^5^ iterations and a run length of 1 × 10^6^ iterations. The number of clusters was estimated using the DeltaK criterion of Evanno et al. (2005). The Pophelper R package v2.3.1 (Francis, 2017) was used to summarise assignment results across independent runs and graphically visualise the results. CLUMPP version 1.1.2 (Jakobsson and Rosenberg, 2007) was used to align clustering outputs from multiple runs of STRUCTURE. LargeKGreedy algorithm (M = 3) was chosen with GREEDY OPTION = 2, to test randomly 20 input orders of runs (REPEATS = 20). Genetic structure was also inferred by factorial correspondence analysis (FCA) of population multiscores using Genetix software v4.05.

A hierarchical analysis of molecular variance (AMOVA, Excoffier et al., 1992), as implemented in the poppr R package with the function poppr.amova(), was performed using all loci. The different hierarchical levels were defined as follows: the uppermost level of genetic structure corresponds to the two main clusters N and S Europe, the second level corresponds to the sub-clusters in each cluster, called here regions, the lowest level corresponds to the sampling locations, called here populations. Differences between hierarchical levels were tested by 999 permutations of the data with the function randtest() (ade4 R package v1.7-22, Thioulouse et al., 2018).

Structure analyses were complemented with a discriminant analysis of principal components (DAPC) implemented in the R package adegenet 2.1 (Jombart et al., 2008 & 2010). This approach is free of assumptions about Hardy-Weinberg equilibrium or linkage disequilibrium and allows clusters of genetically related individuals to be identified and described. DAPC analyses were carried out twice. In the first analysis, the find.clusters() function was used to determine the number of groups (K) *de novo*. Up to 24 groups (K) were tested, which corresponds to the number of sampling sites. Optimal K was selected as that with the lowest BIC value (Bayesian Information Criterion). In the second analysis, in order to investigate more precisely the potential source populations of farmed individuals, the dataset was split into two parts: one containing the 599 wild individuals, used as “training dataset”, and one containing the 90 farmed individuals, used as supplementary individuals. Sampling locations were used as *a priori* groups. The assignment of supplementary individuals to groups using the predict.dapc() function was then examined. The optimal number of principal components (PCs) to use in the DAPC was determined using the cross-validation method (xvalDapc() command) and corresponded to 80 (number of PCs with lowest root mean squared error).

Finally, individual assignment tests were performed to assign farmed sporophytes to the population they have the highest probability of belonging to, using the software GeneClass2 (Piry et al., 2004). The “assign/exclude population as origin of individuals” option was used with natural populations of Saint-Brieuc (France), Atlantic Bridge (Scotland) and Frøya (Norway) as reference populations. These populations corresponded to the populations where the parents of the farmed sporophytes might come from. The Bayesian method of Rannala and Mountain (1997) was selected to compute the likelihood of a multilocus genotype occurring in a population. The relative assignment scores of the individuals were examined at the first and second ranks. For an individual *i* in a population *l*, the score was defined as follows: 𝑠𝑐𝑜𝑟𝑒_𝑖,𝑙_ = 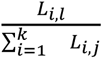 with *L_i,l_* the likelihood value of the individual *i* in the population *l*.

## RESULTS

A total of 221 alleles were identified across the 27 sampled sites (Figure 1a, Table 1) resulting in 688 unique multilocus genotypes. The mean number of alleles per locus was 10.52 ± 7.93 (ranging from 3 to 33). There was 3.82 % missing data across the final data set. Microchecker analysis found no evidence of stuttering error or large allele dropout but did suggest the presence of null alleles in 14 of the 21 loci. For 9 loci (Sacl-11, Sacl-21, Sacl-37, Sacl-41, Sacl-54, Sacl-78, SLN32, SLN320, SLN510), null alleles were not consistent across populations, affecting no more than two populations. For the remaining 5 loci (Sacl-56, SLN34, SLN35, SLN36, SLN54), null alleles were present in 3 to 7 populations, but the average frequency of null alleles remained low (< 0.20, Dakin & Avise, 2004). The average null allele frequency per locus ranged from 0.003 ± 0.013 for Sacl-21 to 0.101 ± 0.134 for SLN34 (see supplementary Table 1 and 2 for populations affected by null alleles and averaged null alleles frequencies per locus).

Genotypic analysis using STRUCTURE and factorial correspondence analysis (FCA) revealed two main clusters: (1) Southern Europe and (2) Northern Europe (Figure 1b, Figure 2). A hierarchical analysis of genetic structure further subdivided the Southern Europe cluster into four sub-clusters and the Northern Europe cluster into two sub-clusters resulting in six sub-clusters : Portuguese and Spanish populations (sub-cluster 1), southern Brittany (sub-cluster 2), northern Brittany and English Channel (sub-cluster 3) Audresselles and Helgoland (sub-cluster 4), Irish and Scottish populations (sub-cluster 5), Swedish and Norwegian populations (sub-cluster 6) (Figure 1c and supplementary Figure 1 for DeltaK plots).

**Fig. 2.**
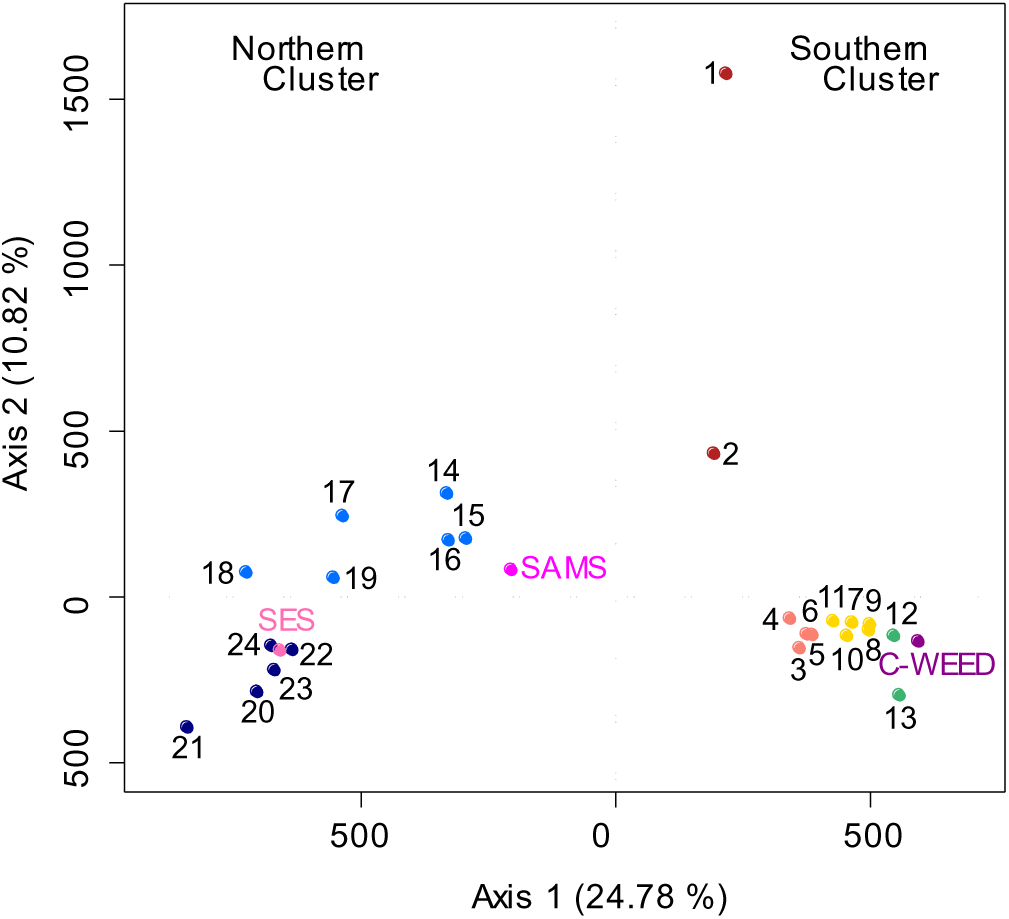
Genetic differentiation of *Saccharina latissima* populations, inferred from factorial correspondence analysis (FCA) of population multivariate scores. Numbers correspond to the sampling sites listed in Figure 1 and Table 1

The separation into two main clusters was also supported by the first DAPC analysis, but subsequent hierarchical levels were not well-supported. The BIC values showed a clear decrease for K=2, but continued to decrease until K=8 and remained more or less stable afterwards (supplementary Figure 2). AMOVA results indicated significant population differentiation at all hierarchical levels: between clusters, between regions within clusters, between populations within regions and within populations (all p values < 0.001, Table 2). Fifty-three percent of the total variance was attributed to differences within populations, while 27 % was due to differences between clusters, whereas differences between populations within regions and between regions within clusters each contributed to 10 % of the total variance.

**Table 2.** Hierarchical analysis of molecular variance (AMOVA). Df: degrees of freedom, SS: sum of squares, MS: mean sum of squares, Est. Covar.: estimated covariance, % Total: percentage of total covariance, Φ-statistics indicate genetic differentiation at different hierarchical levels: between clusters (ɸ_CT_), between regions within clusters (ɸ_RC_), between populations within regions (ɸ_PR_), and within populations (ɸ_PT_). Alter.: alternative hypothesis, p-value: significance of 999 permutation tests

| Source of variation | Df | SS | MS | Est. Covar. | % Total | $\phi$ | Alter. | P-value |
| --- | --- | --- | --- | --- | --- | --- | --- | --- |
| Between clusters | 1 | 730.36 | 730.36 | 2.085 | 26.69 | $\phi_{CT} = 0.267$ | greater | 0.001 |
| Between regions within clusters | 4 | 386.43 | 96.61 | 0.772 | 9.88 | $\phi_{RC} = 0.135$ | greater | 0.001 |
| Between populations within regions | 18 | 433.23 | 24.07 | 0.824 | 10.54 | $\phi_{PR} = 0.166$ | greater | 0.001 |
| Within populations | 575 | 2376.09 | 4.13 | 4.132 | 52.89 | $\phi_{PT} = 0.471$ | less | 0.001 |
| Total | 598 | 3926.12 | 6.56 | 7.813 | 100 |  |  | 0.001 |

Null alleles had some effect on the genetic structure: the ENA method gave slightly, but significantly, lower *F*_st_ values (average *F*_st_ with ENA = 0.268 ± 0.132) than those obtained without ENA correction for the presence of null alleles (average *F*_st_ without ENA = 0.274 ± 0.133; paired t = 19.253, p < 2.2e-16). However, the most and least differentiated pairwise comparisons and the main pattern of differentiation between northern and southern populations persisted with the ENA correction (supplementary Table 3).

The highest genetic diversity was found in the northern cluster, with standardised allelic richness (Â) = 8.67, standardised private allelic richness (PÂ) = 54.57 ± 1.79 and expected heterozygosity (He) = 0.48 (Figure 1). Within this group, sub-cluster 5 (Ireland and Scotland populations) exhibited the highest genetic diversity, with Â = 6.17 ± 0.22, PÂ = 21.75 ± 3.13, He = 0.47. Sub-cluster 3, corresponding to northern Brittany and English Channel populations, also show relatively high genetic diversity (third highest expected heterozygosity and private allelic richness, second highest allelic richness). In contrast, sub-cluster 1, representing Portuguese and Spanish populations, had lower genetic diversity in terms of expected heterozygosity (He = 0.27) and number of different alleles (Â = 4.1), but exhibited the second-highest private allelic richness (PÂ = 11.14 ± 1.51).

Among natural populations, standardised allelic richness (Â) ranged from 1.8 ± 0.08 in Helgoland (Germany) to 4.21 ± 0.18 in Ventry Bay (Ireland) (Table 1). In the farmed populations, Â ranged from 1.94 ± 0.11 (SES farm, Norway) to 2.36 ± 0.15 (C-WEED farm, France). Standardised private allelic richness (PÂ) varied from 0.01 ± 0.11 in Timret (Sweden) to 7.7 ± 2.12 in Ventry Bay (Ireland) within natural populations and between 0.12 ± 0.34 (SES farm, Norway) and 0.47 ± 0.54 (SAMS farm, Scotland) for cultivated populations. Expected heterozygosity (He) ranged from 0.155 in Helgoland (Germany) to 0.461 in Ventry Bay (Ireland) within natural populations and from 0.229 in SES farm (Norway) to 0.371 in SAMS farm (Scotland) for cultivated populations. Significant positive F_is_ values were obtained for 12 of the 24 natural populations and for two of the three farms. Population selfing rate (*s*), varied from 0 to 0.209 within natural populations and from 0.215 to 0.405 within farms. The estimator of selfing *s* was lower in the SES farm, than in the C-WEED and SAMS farms. All three farms, and five natural populations (Ría de Muros y Noya in Spain, Concarneau in Southern Brittany, France, and Saint-Malo in Northern Brittany, France, Shieldaig in Scotland and Timret in Sweden), showed significant correlation in heterozygosities across loci. Significant linkage disequilibria were only found within the natural population of Porthallow (England) and within all the three farms.

All the three methods used to assign the cultivated sporophytes to natural populations (STRUCTURE, DAPC and GenClass2) indicated no clear genetic separation between farmed and wild populations. The Bayesian analysis in the STRUCTURE revealed that individuals from the C-WEED farm were assigned to the southern cluster (sub-cluster 3), while those from the SAMS or SES farms were assigned to the northern cluster (sub-cluster 5 and 6, respectively) (Figure 1). DAPC analysis, using wild sporophytes as a training dataset and farmed sporophytes as test individuals, revealed that the percentage of sporophytes correctly assigned to their parental population was higher for the C-WEED farm (91.43 %), than for the SAMS (76 %) and SES farms (66.67 %) (Figure 3 and 4).

**Fig. 3.**
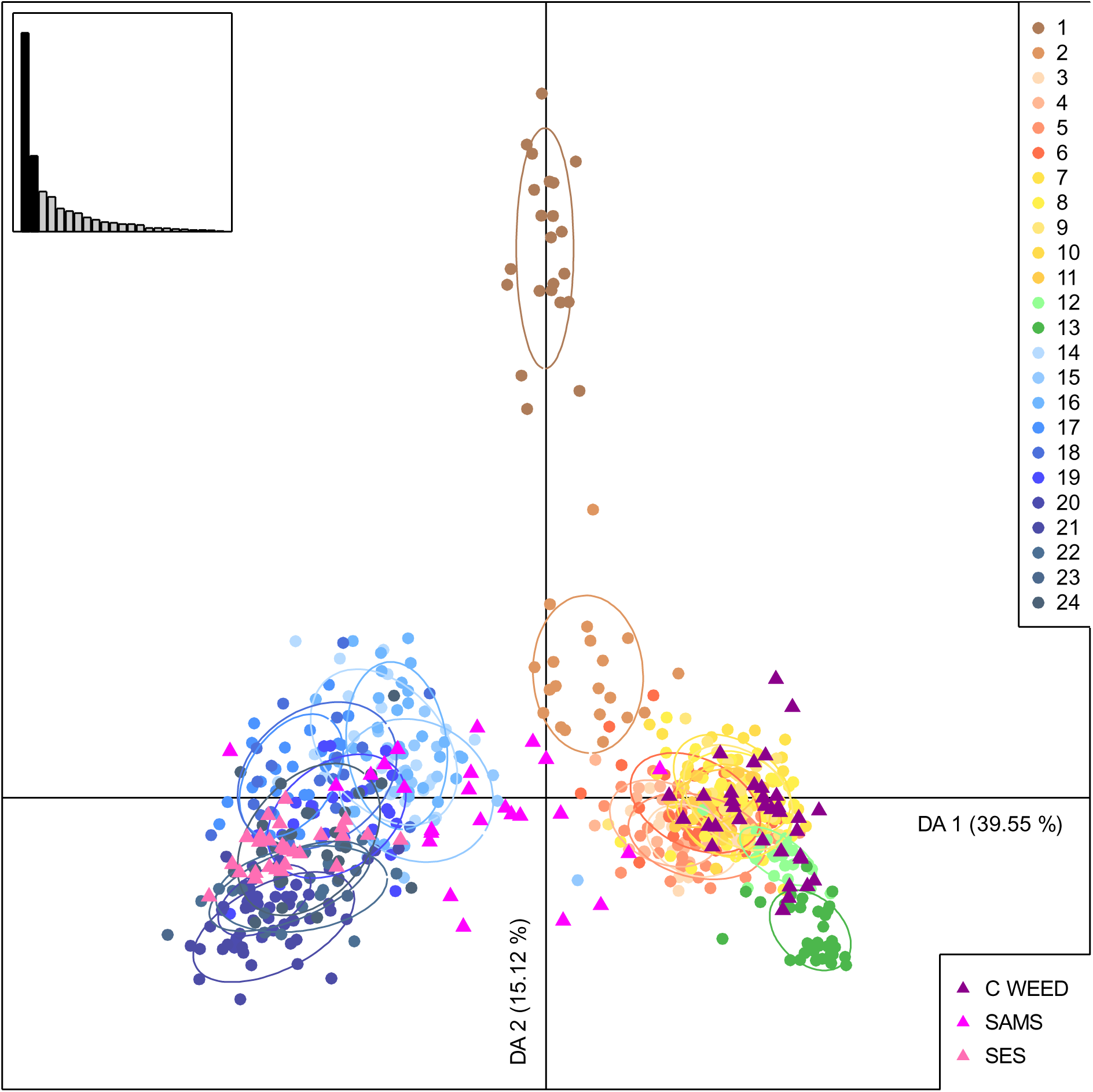
Discriminant Analysis of Principal Components (DAPC) with supplementary individuals. Training dataset individuals are represented by dots, and population groups by inertia ellipses. Individuals are colored according to STRUCTURE and FCA results. Eigenvalues for the analysis are shown in the inset. Supplementary individuals from the farmed populations are indicated by triangles. Numbers refer to sites listed in Figure 1 and Table 1

**Fig. 4.**
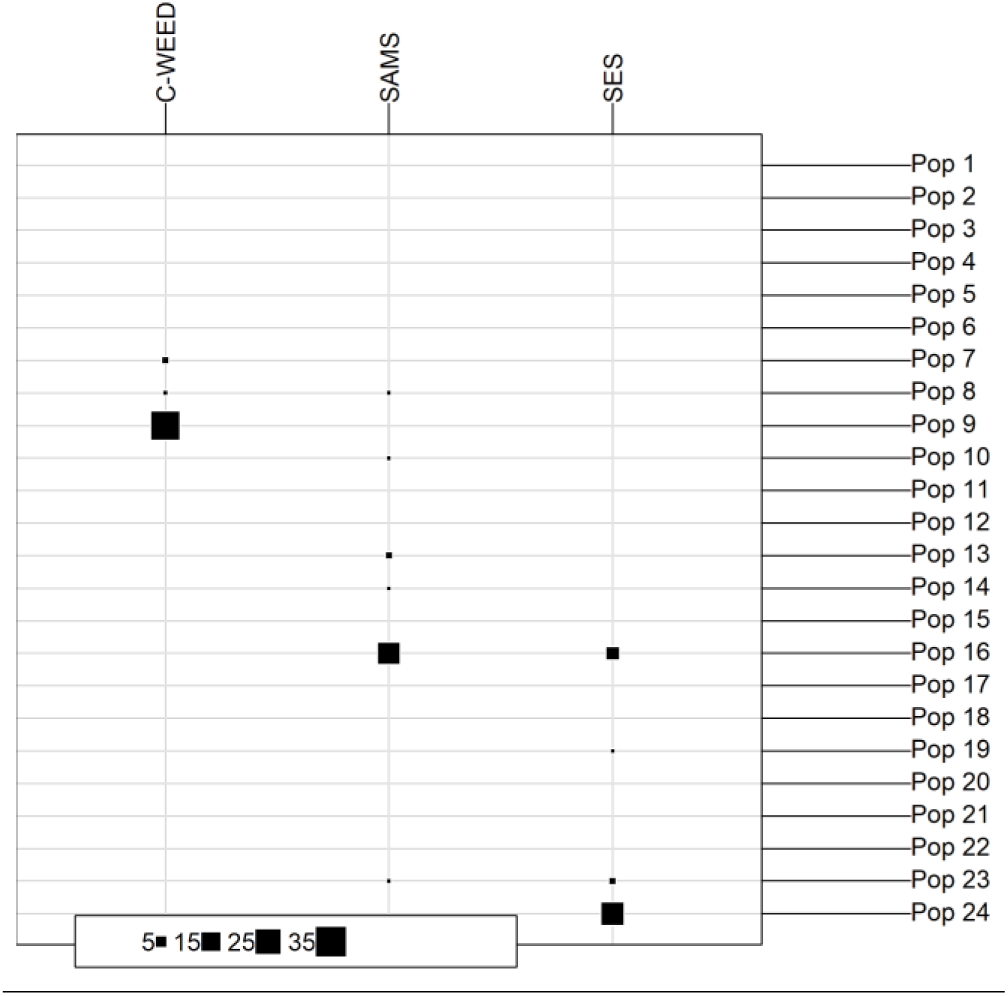
Assignment of farmed sporophytes (supplementary individuals) to their potential natural populations using DAPC. Columns represent the farmed populations and rows the natural populations. Numbers refer to sites as listed in Figure 1 and Table 1

GeneClass2 assignment tests support this pattern, with all individuals from the C-WEED and SAMS farms correctly assigned to their populations of origin, achieving very high assignment scores (100 % ± 0 for C-WEED and 99.9 % ± 0.04 for the SAMS). In contrast, only 63.3 % of the individuals from the SES farm were correctly assigned to their population of origin with an average assignment score of 64.5 % (Figure 5 and supplementary Table 3). Regardless of the method used, assignment scores were consistently higher for the C-WEED farm compared to the SAMS and SES farms.

**Fig. 5.**
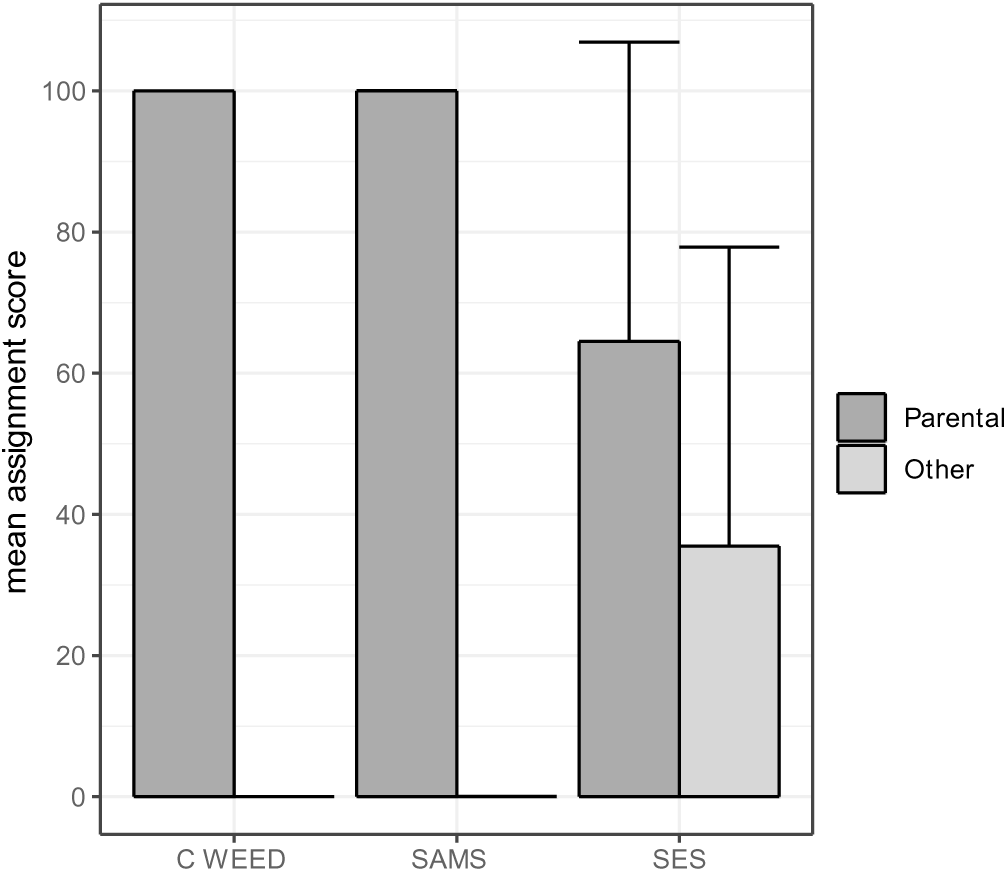
Mean assignment score of the farmed sporophytes from the C-WEED, SAMS and SES farms to their parental populations, as estimated using GeneClass2 software. Error bars represent standard deviation

## DISCUSSION

In comparison with a previous study that analysed the whole range of distribution of the *S. latissima* species complex in the North Atlantic and North-East Pacific (Neiva et al., 2018), the present study focuses on the genetic structure of *S. latissima* along the European coasts. Our analyses showed different levels of hierarchical genetic structure of *S. latissima* in the North-East Atlantic associated with contrasted patterns of genetic diversity reflecting the historical processes that occurred after the Last Glacial Maximum (LGM) and current selection processes possibly related to climate change. Despite a situation of relative high connectivity between wild populations within regions (populations separated by up to hundreds of kilometers appeared to be connected by gene flow), the cultivated genotypes were fairly correctly assigned to their population of origin in three cultivation sites in Europe, and we were even able to detect an undeclared mixture in the provenance of the parental sporophytes for one of the farms. These findings are discussed below in relation to previous studies on this species and other kelps, with a particular focus on the decline of kelp forests due to climate change and other stressors. Additionally, the discussion considers the context of the development of the European seaweed aquaculture sector, particularly concerning strain traceability.

### Two well separated genetic clusters of *S. latissima* in Europe

The results of the present study confirm strong genetic differentiation between Northern and Southern Europe and refine the hierarchical genetic structure of the species in Europe as suggested in previous studies (Guzinski et al., 2016; 2020; Møller Nielsen et al., 2016; Luttikhuizen et al., 2018; Mooney et al., 2018; reviewed in Diehl et al., 2024). The occurrence of two well separated genetic clusters of *S. latissima* in Europe has not only been confirmed, but the boundary between these two groups has been significantly narrowed compared with the previous study by Neiva et al. (2018), which differentiated the southern populations of Galicia and Brittany from the northern populations located north of Norway. In our study, the southern boundary of the northern cluster extends to Southern Ireland, and the northern boundary of the southern cluster to the English Channel and the southern part of the North sea.

The presence of distinct genetic clusters in Europe, arising from historical processes implying recolonisation after the Last Glacial Maximum and originating from various refugia has been documented for many European coastal species (see review for marine benthic organisms Maggs et al., 2008, for seaweed Neiva et al., 2016 and for Atlantic kelps Assis et al., 2018). The boundary between the two genetic clusters identified in our study closely resembles that of the amphi-Atlantic kelp species *Laminaria digitata* (Neiva et al., 2020; Reynes et al., 2024) suggesting recolonisation from Southern Europe (northwest Iberia and Celtic Sea) on one hand, and from Northern Europe (due to the persistence of kelps during glaciation at high latitude further than northern Norway), see references above. Past range dynamics was probably very different between northern and southern Europe explaining the observed difference in genetic diversity and structure between the two genetic clusters.

### Higher genetic diversity in the Northern European cluster of *S. latissima*: implications for climate change resilience

The northern cluster of *S. latissima* is characterised by a higher genetic diversity and allelic richness but a simpler genetic structure than the southern one. Indeed, the southern cluster groups four genetic sub-clusters while the northern one is composed of only two genetic sub-clusters. The vulnerability of *S. latissima* to high temperatures has probably impacted its range dynamics in Southern Europe during the last LGM leading to reduced population sizes and limited gene flow. This pattern of low genetic diversity within population combined with high genetic differentiation among populations is characteristic of low-latitude rear-edge populations (Hampe and Petit, 2005) and has been reported in many kelp species across southern Europe, including *S. polyschides* (Assis et al., 2016), *L. digitata* (Neiva et al., 2020) and *S. latissima* (Guzinski et al., 2020).

In the context of climate change, the contemporary genetic pools of the different regions in Southern Europe are threatened by extinction in the near future, as projected for most kelp populations located at lower latitudes (Assis et al., 2018; Nimbs et al., 2023; Assis et al., 2024). For *S. latissima,* populations exhibiting lower genetic diversity have predominantly been reported in areas where a decline of species abundance have been observed over the last two decades. In the strait of Dover, Portugal and northern Spain, the loss of populations can be partly attributed to high temperature events (Filbee-Dexter et al., 2020; Nepper-Davidsen et al., 2020). However, changes in kelp distributions in response to shifts in sea temperature may be influenced by acclimation and local adaptations and not only by maladaptation resulting from genetic drift. A recent study in the kelp *L. digitata* in the North-East Atlantic has shown the promise of temporal genomics to gain deeper insights into the contemporary responses of this fundamental marine species to rapid environmental changes (Reynes et al., 2024). By employing ddRADseq on four populations, this temporal approach allowed the effects of ongoing selection relative to genetic drift on changes in allele frequency to be deciphered.

In the northern cluster, wild populations of *S. latissima* from Ireland and Scotland exhibited the highest levels of genetic diversity, consistent with long-term persistence at large effective population sizes. This finding supports the hypothesis that high-latitude North Atlantic and Arctic refugia have persisted through glaciation cycles (Assis et al., 2018; Bringloe et al., 2022). In contrast to the southern rear-edge, the gene pools of northern populations contain the variability necessary for potential adaptation through natural selection under future conditions (Wernberg al., 2010; 2018). These northern populations are expected to demonstrate greater resilience to future climate change compared to populations at the southern rear-edge. Furthermore, they may offer a valuable opportunity for aquaculture to enhance resilience to climate change (Veenhof et al., 2024).

### Not all farmed genotypes could be assigned to their source population

The genetic structure among wild populations within clusters was minimal even across distances ranging from 40 km to over a thousand kilometers. This was particularly evident in the Northern European cluster, where only two genetic groups were identified. Despite this low level of differentiation, our results demonstrated that all farmed genotypes were assigned to their genetic cluster of origin. However, it is important to note that not all cultivated genotypes could be definitively assigned to their source population. In the southern genetic cluster, farmed individuals were not only accurately assigned to their wild source populations but were also distinguishable from other populations within the same genetic group, even when those populations were geographically closer than the source population. Conversely, in the northern genetic cluster, where the genetic structure is less pronounced, the majority of cultivated individuals were still assigned to the population from which the parents originated, although a small number of individuals was also assigned to more distant populations.

The assignment scores varied among the three farms, likely reflecting differences in cultivation practices. The C-WEED farm, located in Brittany less than 10 km from the wild population of Saint-Malo (pop 10), was assigned to the wild population of Saint-Brieuc (pop 9), which is over 80 km away. This assignment is consistent with the fact that the parental sporophytes used to seed the farm were collected from Saint-Brieuc. C-WEED is a family-run farm that cultivates and harvests seaweed primarily for the food industry, as well as for cosmetics, to produce high-value products. They commercialize their own line of dried edible seaweed and hold an organic certification. Their production methods adhere to the principles of agrobiology and sustainable resource management, ensuring maximum traceability of their seaweed.

The SAMS farm is an experimental farm which aims to provide material for research studies. The biological material utilized from the SAMS farm in this study was not initially intended for investigating gene flow. The origin of the parental sporophytes was not relevant for the ongoing experiment, and only a few individuals from two different localities were used as parental broodstock. Consequently, it was not feasible to genotype the true parental population, and the population used for the comparison did not correspond to the one that supplied the parental material for the cultivation lines, which accounts for the lower assignment scores observed.

In contrast, the situation at the SES farm in Norway is more complex, as only 63% of the individuals were assigned to their population of origins, suggesting potential contamination from another nearby, differentiated population. The SES farm is one of the largest seaweed farms in Europe, supplying raw material to the food industry and other markets. While the parental individuals are believed to primarily originated mainly from the wild population of Frøya, discussions with the farmers revealed that the seedlings were indeed a mix from two locations, with some individuals possibly sourced from the fjord of Trondheim, located 80 kilometers away from the farm.

Our study highlights the effectiveness of microsatellites in facilitating population genetics analyses and in providing support for breeding, selection and conservation programs even when compared to more recent but more expensive markers such as SNPs or sequence data (Mauger et al., 2023). The 21 markers enabled the differentiation of genetic resources and strains, allowing for the detection of gene flow between farms and wild populations. Similar findings have been reported in Portugal, where microsatellite markers successfully confirmed the genetic assignment of aquaculture breeder samples to their natural source populations (E. Serrão comm. pers.). However, the use of denser genomic markers such as single nucleotide polymorphisms (SNPs), showed more power than microsatellites for identifying fine grained patterns (Guzinski et al., 2018; Bratelund et al., 2024).

### Farms exhibit reduced genetic diversity and increased inbreeding compared to their source populations

Domestication typically leads to reduction in allelic variation and genetic diversity, resulting in genetic erosion, as observed in terrestrial crops (Diamond, 2002). In our study, genetic diversity within the farmed populations was consistently lower than in their source populations, as evidenced by lower expected heterozygosity and allelic richness. In the three farms, cultivation lines were seeded directly with zoospores produced by a relatively small number of sporophytes sourced from wild populations. The broodstock is notably limited, with numbers of sporophytes ranging from approximately a dozen at the SAMS farm, to 200-300 at the SES farm, and around 30 to 40 at the C-WEED farm. This limited genetic pool constitutes only a small fraction of the genetic diversity found in wild populations. Consequently, the cultivated populations exhibit reduced genetic diversity and increased inbreeding, as a result of reduced effective population sizes. Our results indicate a shift in the mating system in farms with increased inbreeding compared to wild populations. The seedling production methods used in farms have led to higher levels of self-fertilisation and crosses between related gametophytes, as a result of a limited number of parental sporophytes being used. A correlation was observed between the level of selfing (*s,* see Table 1) and the amount of parental material used to seed the lines on the farms. To reduce the effect of inbreeding depression in this allogamous species, it is recommended that farmers use a larger number of parental sporophytes.

Such loss of diversity has also been documented in other cultivated allogamous seaweeds with large population sizes when seedlings are produced from a reduced number of parents compared to the natural population (*Saccharina japonica*: Shan et al., 2011; Liu et al., 2012, Hu et al., 2021; *Ulva prolifera*: Huh et al., 2004); *Porphyra yezoensis*: Kodaka, Akiyama and Fuji, 1988). In contrast to what was observed in the outcrossing *S. latissima*, a shift in a mating system of the cultivated kelp autogamous *Undaria pinnatifida*, has been reported, with reduced levels of inbreeding in farmed populations compared to wild populations in Brittany (Guzinski et al., 2018) and Korea (Graf et al., 2021). Breeding practices using artificial mixtures have counteracted the naturally high rates of self-fertilisation characteristic of this autogamous species. However, the genetic diversity observed in farms is reduced compared to wild populations in Brittany while this reduction is not evident in Korea where *U. pinnatifida* is cultivated on a large scale in extensive farms, which contrast markedly with the farming practices employed in Brittany. In Korea, breeding occurs in large pools where multiple sporangia from both cultivated and wild individuals are combined, thereby facilitating recruitment on culture ropes. Consequently, employing a high number of parental sporophytes for seedling the lines, as practiced in Korea, is crucial for minimising the loss of genetic diversity in cultivated populations.

### Implications for strain traceability and aquaculture practices

As pointed out recently by Bemmels et al. (2024) for the management of the giant kelp *Macrocystis pyrifera* populations along the North-West American coast, guidelines are essential to monitor the movement of genetic material for kelp aquaculture and restoration. While European regulations do not specifically address seaweed, they prohibit the use of foreign strains and non-local genotypes in aquaculture practices (Barbier et al., 2019). Farmers currently use parental sporophytes directly sourced from local wild populations, with the definition of local population remaining subjective. Our findings confirm that wild populations of *S. latissima* exhibit significant genetic structure. These populations are often locally adapted to their specific environments and display unique morphological and phenological traits. Given these considerations, we recommend the use of the genetic clusters identified in this study to define the geographic origin of parental sporophytes in relation to the location of the aquaculture farm. Moreover, translocation of cultivated material outside of its genetic cluster of origin should be avoided to prevent the introduction of genotypes that could disrupt local wild populations.

The majority of nurseries depend on collecting wild parental sporophytes to produce seeds. However, some wild populations are at risk of extinction due to climate change and other stressors (e.g., populations in Audresselles, Helgoland, or Portugal). Additionally, sourcing from wild populations limits the ability to control for specific desirable traits. The unpredictable quality and quantity of seeds can lead to delays in farm deployment and/or reduced yields, ultimately affecting the entire seaweed farming value chain. Hence, it is essential to create a collection of both wild and cultivated strains of *S. latissima* to preserve wild genetic diversity over the long term, maintain the best cultivars, and ensure a variety of strains for breeding and selection programs (Hofmann et al., in review). However, this ex-situ conservation strategy does not take into account the dynamics of genetic diversity in response to environmental changes and questions about what *in situ* management of genetic resources could be implemented. It is therefore important in aquaculture not only to use local cultivars, but also to maintain a large effective population size and frequently introduce new genotypes from wild type brood stock or diverse stocks of gametophytes from the same locality (genetic region) to ensure high genetic diversity among cultivars (Barbier et al., 2019; Goecke et al., 2020; Graf et al., 2021; Hu et al., 2021; 2024).

## Declarations

### Funding

This work was supported by the European Union’s Horizon 2020 research and innovation programme under grant agreement No. 727892 (GENIALG). This output reflects only the author’s view, and the European Union cannot be held responsible for any use that may be made of the information contained therein. CNRS and Sorbonne Université core fundings to the International Research Laboratory (IRL) 3614, EBEA Evolutionary Biology and Ecology of Algae and UMR 8227 are also acknowledged.

### Competing interests

The authors declare no competing interests.

### Availability of data and material

Microsatellite data used for genetic analyses have been deposited in Data.InDoRES repository at https://www.indores.fr/.

### Code availability

Not applicable.

### Authors’ contributions

L. Jaugeon: sampling, DNA extraction and genotyping, bioinformatics analysis, analysis of molecular data, led manuscript drafting and editing; C. Destombe: sampling, original concept, led manuscript drafting and editing; P. Ruggeri: DNA extraction and genotyping and editing manuscript; S. Mauger: genotyping and editing manuscript; J. Coudret: sampling and editing manuscript; J.M. Cock: funding and editing manuscript; P. Potin: original concept, funding and led manuscript drafting and editing; M. Valero: original concept, funding and supervision, sampling, drafting and editing manuscript. All authors read and approved the final manuscript.

## Acknowledgments

We are grateful to all the GENIALG partners who participated in sampling and providing us either natural or cultivated samples of *Saccharina latissima*. The Scottish Association for Marine Science (SAMS), National University of Galway (NUIG), Centro Interdisciplinar de Investigação Marinha e Ambiental (CIIMAR), Seaweed Energy Solutions (SES) and C-Weed Aquaculture SAS. We also thank the Service Mer & Observation of Roscoff and our colleagues from IRL EBEA 3614 team for sampling. In addition, we are grateful for the contribution of Wouter Visch (University of Gothenburg - Tjärnö), Inka Bartsch (Alfred Wegener Institute Helmholtz Centre - Bremerhaven), Akira Peters (BEZHIN ROSKO) who arranged access to populations of Sweden, Germany and Pays de la Loire (France), respectively. We warmly thank François Gevaert (University of Lille, Wimereux Marine Station) for his help in finding the vanishing populations of *S. latissima* in Audresselles. We are also grateful to the Genomer Platform, EMBRC-France partner core facility and member of Biogenouest, the Roscoff Aquarium Services (RAS) team and the Service Cultures d’Algues at the Station Biologique de Roscoff for their technical support.

## Notes

### Competing Interest Statement

The authors have declared no competing interest.

