## Supplementary figures for "Wild and farmed *Saccharina latissima* in Europe: genetic insights for sustainable cultivation, traceability and environmental challenges"

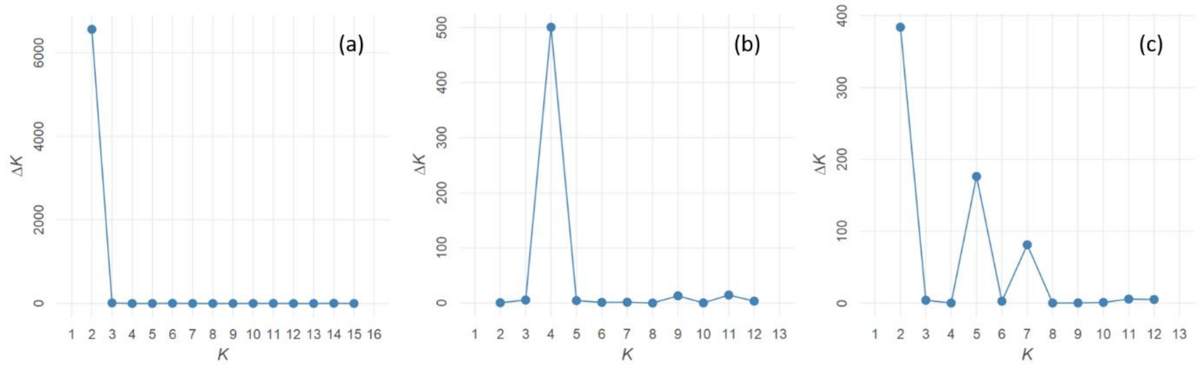


**Supplementary Figure 1:** DeltaK values (DeltaK, a measure of the rate of change in the STRUCTURE likelihood function) as a function of K, the number of putative populations. Graphs (a), (b) and (c) show the results for the entire dataset, within southern clade and within northern clade, respectively.

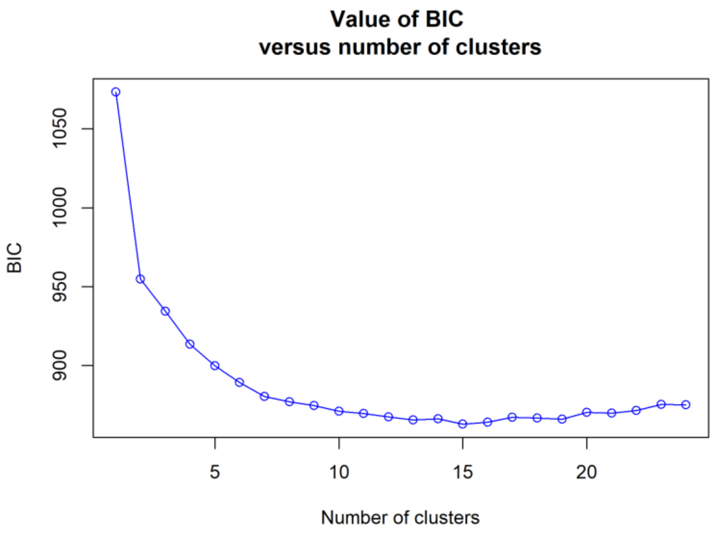


**Supplementary Figure 2:** Bayesian information criterion (BIC) plot from the find.clusters() function
