## Supplementary tables for "Wild and farmed *Saccharina latissima* in Europe: genetic insights for sustainable cultivation, traceability and environmental challenges"

**Supplementary table 1:** Loci potentially affected by null alleles. Results of the Microchecker v. 2.2.3 analysis for each locus and population. See table 1 and Figure 1 for the names of populations.

| Locus/Pop | 1 | 2 | 3 | 4 | 5 | 6 | 7 | 8 | 9 | 10 | 11 | 12 | 13 | 14 | 15 | 16 | 17 | 18 | 19 | 20 | 21 | 22 | 23 | 24 | 25 | 26 | 27 |
|---|---|---|---|---|---|---|---|---|---|---|---|---|---|---|---|---|---|---|---|---|---|---|---|---|---|---|---|
| Sacl-11 | no | no | no | no | no | no | no | no | no | no | no | no | no | no | no | no | no | no | **yes** | no | no | no | no | no | no | no | no |
| Sacl-37 | no | no | no | no | no | no | no | no | no | no | no | no | no | no | no | no | no | no | **yes** | no | **yes** | no | no | no | no | no | no |
| Sacl-54 | no | no | no | **yes** | no | no | no | no | no | no | no | no | no | no | **yes** | no | no | no | no | no | no | no | no | no | no | no | no |
| Sacl-65 | no | no | no | no | no | no | no | no | no | no | no | no | no | no | no | no | no | no | no | no | no | no | no | no | no | no | no |
| Sacl-95 | no | no | no | no | no | no | no | no | no | no | no | no | no | no | no | no | no | no | no | no | no | no | no | no | no | no | no |
| Sacl-13 | no | no | no | no | no | no | no | no | no | no | no | no | no | no | no | no | no | no | no | no | no | no | no | no | no | no | no |
| Sacl-21 | no | no | **yes** | no | no | no | no | no | no | no | no | no | no | no | no | no | no | no | no | no | no | no | no | no | no | no | no |
| Sacl-33 | no | no | no | no | no | no | no | no | no | no | no | no | no | no | no | no | no | no | no | no | no | no | no | no | no | no | no |
| Sacl-41 | no | no | no | no | no | no | no | no | no | no | no | no | no | no | no | no | **yes** | no | no | no | no | no | no | no | no | no | no |
| Sacl-56 | no | no | no | no | no | no | no | no | no | no | no | no | no | no | **yes** | no | **yes** | **yes** | no | no | no | no | no | no | no | no | no |
| Sacl-60 | no | no | no | no | no | no | no | no | no | no | no | no | no | no | no | no | no | no | no | no | no | no | no | no | no | no | no |
| Sacl-78 | no | no | no | no | no | no | no | no | no | no | no | no | no | no | no | no | no | no | **yes** | no | no | no | no | no | no | no | no |
| Sacl-88 | no | no | no | no | no | no | no | no | no | no | no | no | no | no | no | **yes** | no | no | no | no | no | no | no | **yes** | no | no | no |
| SLN319 | no | no | no | no | no | no | no | no | no | no | no | no | no | no | no | no | no | no | no | no | no | no | no | no | no | no | no |
| SLN320 | no | no | no | no | no | no | **yes** | no | no | no | no | no | no | no | no | no | no | no | no | no | no | no | no | no | no | no | no |
| SLN34 | no | no | no | no | no | no | no | no | no | no | no | no | no | no | no | no | **yes** | no | **yes** | **yes** | **yes** | **yes** | **yes** | **yes** | no | no | no |
| SLN35 | no | no | no | no | no | no | no | no | no | **yes** | no | no | no | no | no | no | no | no | **yes** | no | **yes** | no | no | **yes** | no | **yes** | no |
| SLN 510 | no | no | no | no | no | no | no | no | no | no | no | no | no | no | no | no | no | no | no | no | no | no | no | **yes** | no | no | no |
| SLN 32 | no | no | no | no | no | no | no | no | no | no | no | no | no | no | no | no | no | no | no | no | no | no | no | no | no | no | **yes** |
| SLN 36 | no | no | no | no | no | no | no | no | no | no | no | no | no | no | no | no | no | no | no | no | no | **yes** | no | **yes** | no | **yes** | **yes** |
| SLN 54 | no | no | no | no | **yes** | no | **yes** | **yes** | no | no | no | no | no | no | no | no | no | no | no | **yes** | **yes** | no | **yes** | no | no | no | no |

**Supplementary table 2:** average frequency of null alleles per locus and associated standard deviation. The frequency of null alleles per locus and per population was estimated according to the EM algorithm (Dempster et al. 1977) using FreeNA software (Chapuis and Estoup, 2007).

| Locus | mean | sd |
|---|---|---|
| Sacl_11 | 0.028 | 0.062 |
| Sacl_37 | 0.027 | 0.063 |
| Sacl_54 | 0.035 | 0.058 |
| Sacl_65 | 0.035 | 0.066 |
| Sacl_95 | 0.025 | 0.041 |
| Sacl_13 | 0.005 | 0.021 |
| Sacl_21 | 0.003 | 0.013 |
| Sacl_33 | 0.005 | 0.023 |
| Sacl_41 | 0.016 | 0.047 |
| Sacl_56 | 0.038 | 0.062 |
| Sacl_60 | 0.013 | 0.032 |
| Sacl_78 | 0.018 | 0.042 |
| Sacl_88 | 0.032 | 0.057 |
| SLN319 | 0.012 | 0.03 |
| SLN320 | 0.018 | 0.035 |
| SLN34 | 0.101 | 0.134 |
| SLN35 | 0.058 | 0.061 |
| SLN510 | 0.013 | 0.025 |
| SLN32 | 0.031 | 0.041 |
| SLN36 | 0.034 | 0.055 |
| SLN54 | 0.047 | 0.076 |

**Supplementary table 3:** Estimating Fst of Weir (1996) for each pair of populations both using and without using the ENA correction described in Chapuis and Estoup (2007). The first matrix corresponds to the values of pairwise Fst without using ENA correction and the second matrix to the values of pairwise Fst using the ENA correction. See table 1 and Figure 1 for the names of populations. The darker the color, the higher the pairwise Fst value.

First matrix: without using ENA

| pop | 1 | 2 | 3 | 4 | 5 | 6 | 7 | 8 | 9 | 10 | 11 | 12 | 13 | 14 | 15 | 16 | 17 | 18 | 19 | 20 | 21 | 22 | 23 | 24 | 25 | 26 |
|---|---|---|---|---|---|---|---|---|---|---|---|---|---|---|---|---|---|---|---|---|---|---|---|---|---|---|
| 2 | 0,23 | | | | | | | | | | | | | | | | | | | | | | | | | |
| 3 | 0,36 | 0,23 | | | | | | | | | | | | | | | | | | | | | | | | |
| 4 | 0,32 | 0,17 | 0,07 | | | | | | | | | | | | | | | | | | | | | | | |
| 5 | 0,32 | 0,21 | 0,03 | 0,06 | | | | | | | | | | | | | | | | | | | | | | |
| 6 | 0,30 | 0,18 | 0,07 | 0,09 | 0,07 | | | | | | | | | | | | | | | | | | | | | |
| 7 | 0,30 | 0,19 | 0,10 | 0,11 | 0,11 | 0,05 | | | | | | | | | | | | | | | | | | | | |
| 8 | 0,31 | 0,21 | 0,10 | 0,13 | 0,11 | 0,07 | 0,01 | | | | | | | | | | | | | | | | | | | |
| 9 | 0,30 | 0,16 | 0,11 | 0,11 | 0,09 | 0,08 | 0,03 | 0,02 | | | | | | | | | | | | | | | | | | |
| 10 | 0,32 | 0,16 | 0,13 | 0,14 | 0,12 | 0,10 | 0,07 | 0,06 | 0,04 | | | | | | | | | | | | | | | | | |
| 11 | 0,37 | 0,19 | 0,14 | 0,12 | 0,11 | 0,11 | 0,07 | 0,09 | 0,07 | 0,10 | | | | | | | | | | | | | | | | |
| 12 | 0,43 | 0,30 | 0,18 | 0,21 | 0,15 | 0,15 | 0,12 | 0,13 | 0,15 | 0,20 | 0,20 | | | | | | | | | | | | | | | |
| 13 | 0,59 | 0,44 | 0,35 | 0,42 | 0,35 | 0,32 | 0,27 | 0,27 | 0,31 | 0,36 | 0,44 | 0,32 | | | | | | | | | | | | | | |
| 14 | 0,30 | 0,17 | 0,23 | 0,22 | 0,26 | 0,22 | 0,23 | 0,27 | 0,24 | 0,23 | 0,24 | 0,33 | 0,44 | | | | | | | | | | | | | |
| 15 | 0,36 | 0,19 | 0,28 | 0,27 | 0,32 | 0,26 | 0,26 | 0,30 | 0,27 | 0,28 | 0,27 | 0,35 | 0,41 | 0,08 | | | | | | | | | | | | |
| 16 | 0,31 | 0,20 | 0,25 | 0,25 | 0,27 | 0,23 | 0,26 | 0,28 | 0,25 | 0,24 | 0,26 | 0,34 | 0,44 | 0,04 | 0,14 | | | | | | | | | | | |
| 17 | 0,34 | 0,21 | 0,32 | 0,30 | 0,34 | 0,30 | 0,32 | 0,35 | 0,31 | 0,30 | 0,30 | 0,40 | 0,51 | 0,10 | 0,14 | 0,13 | | | | | | | | | | |
| 18 | 0,41 | 0,28 | 0,34 | 0,33 | 0,37 | 0,31 | 0,34 | 0,37 | 0,34 | 0,33 | 0,34 | 0,42 | 0,51 | 0,13 | 0,19 | 0,11 | 0,11 | | | | | | | | | |
| 19 | 0,39 | 0,24 | 0,30 | 0,30 | 0,34 | 0,29 | 0,32 | 0,35 | 0,31 | 0,31 | 0,31 | 0,40 | 0,50 | 0,08 | 0,12 | 0,10 | 0,09 | 0,07 | | | | | | | | |
| 20 | 0,49 | 0,33 | 0,38 | 0,39 | 0,42 | 0,35 | 0,37 | 0,39 | 0,38 | 0,36 | 0,40 | 0,48 | 0,52 | 0,18 | 0,20 | 0,18 | 0,19 | 0,17 | 0,17 | | | | | | | |
| 21 | 0,52 | 0,37 | 0,42 | 0,44 | 0,45 | 0,39 | 0,41 | 0,43 | 0,42 | 0,41 | 0,44 | 0,51 | 0,55 | 0,25 | 0,27 | 0,23 | 0,26 | 0,20 | 0,23 | 0,08 | | | | | | |
| 22 | 0,50 | 0,31 | 0,39 | 0,38 | 0,41 | 0,34 | 0,37 | 0,39 | 0,38 | 0,36 | 0,41 | 0,48 | 0,57 | 0,14 | 0,19 | 0,16 | 0,17 | 0,16 | 0,14 | 0,12 | 0,23 | | | | | |
| 23 | 0,51 | 0,32 | 0,38 | 0,39 | 0,42 | 0,33 | 0,36 | 0,38 | 0,38 | 0,36 | 0,41 | 0,48 | 0,55 | 0,11 | 0,16 | 0,11 | 0,16 | 0,11 | 0,10 | 0,06 | 0,14 | 0,06 | | | | |
| 24 | 0,41 | 0,26 | 0,32 | 0,32 | 0,36 | 0,29 | 0,32 | 0,34 | 0,32 | 0,31 | 0,33 | 0,40 | 0,47 | 0,12 | 0,16 | 0,11 | 0,13 | 0,09 | 0,11 | 0,09 | 0,11 | 0,10 | 0,06 | | | |
| 25 | 0,38 | 0,27 | 0,21 | 0,20 | 0,16 | 0,15 | 0,10 | 0,09 | 0,06 | 0,13 | 0,14 | 0,20 | 0,40 | 0,33 | 0,36 | 0,34 | 0,39 | 0,42 | 0,39 | 0,47 | 0,50 | 0,47 | 0,47 | 0,40 | | |
| 26 | 0,38 | 0,23 | 0,32 | 0,32 | 0,32 | 0,27 | 0,29 | 0,31 | 0,28 | 0,25 | 0,29 | 0,37 | 0,43 | 0,15 | 0,20 | 0,10 | 0,19 | 0,20 | 0,19 | 0,23 | 0,27 | 0,22 | 0,20 | 0,18 | 0,37 | |
| 27 | 0,58 | 0,43 | 0,48 | 0,49 | 0,49 | 0,44 | 0,47 | 0,48 | 0,48 | 0,47 | 0,51 | 0,57 | 0,65 | 0,21 | 0,32 | 0,16 | 0,27 | 0,22 | 0,20 | 0,29 | 0,32 | 0,32 | 0,23 | 0,16 | 0,55 | 0,30 |

Second matrix: using ENA

| pop | 1 | 2 | 3 | 4 | 5 | 6 | 7 | 8 | 9 | 10 | 11 | 12 | 13 | 14 | 15 | 16 | 17 | 18 | 19 | 20 | 21 | 22 | 23 | 24 | 25 | 26 |
|---|---|---|---|---|---|---|---|---|---|---|---|---|---|---|---|---|---|---|---|---|---|---|---|---|---|---|
| 2 | 0,22 | | | | | | | | | | | | | | | | | | | | | | | | | |
| 3 | 0,35 | 0,21 | | | | | | | | | | | | | | | | | | | | | | | | |
| 4 | 0,31 | 0,16 | 0,08 | | | | | | | | | | | | | | | | | | | | | | | |
| 5 | 0,32 | 0,20 | 0,04 | 0,06 | | | | | | | | | | | | | | | | | | | | | | |
| 6 | 0,30 | 0,18 | 0,07 | 0,09 | 0,08 | | | | | | | | | | | | | | | | | | | | | |
| 7 | 0,29 | 0,18 | 0,10 | 0,11 | 0,11 | 0,05 | | | | | | | | | | | | | | | | | | | | |
| 8 | 0,31 | 0,21 | 0,11 | 0,13 | 0,11 | 0,07 | 0,01 | | | | | | | | | | | | | | | | | | | |
| 9 | 0,30 | 0,15 | 0,11 | 0,11 | 0,09 | 0,08 | 0,03 | 0,02 | | | | | | | | | | | | | | | | | | |
| 10 | 0,31 | 0,15 | 0,13 | 0,13 | 0,11 | 0,10 | 0,07 | 0,06 | 0,04 | | | | | | | | | | | | | | | | | |
| 11 | 0,35 | 0,17 | 0,14 | 0,12 | 0,11 | 0,11 | 0,08 | 0,10 | 0,07 | 0,09 | | | | | | | | | | | | | | | | |
| 12 | 0,43 | 0,30 | 0,18 | 0,21 | 0,15 | 0,15 | 0,12 | 0,13 | 0,15 | 0,20 | 0,20 | | | | | | | | | | | | | | | |
| 13 | 0,59 | 0,44 | 0,35 | 0,42 | 0,34 | 0,32 | 0,27 | 0,26 | 0,31 | 0,36 | 0,43 | 0,32 | | | | | | | | | | | | | | |
| 14 | 0,30 | 0,17 | 0,23 | 0,22 | 0,26 | 0,22 | 0,23 | 0,27 | 0,24 | 0,23 | 0,24 | 0,33 | 0,44 | | | | | | | | | | | | | |
| 15 | 0,35 | 0,18 | 0,26 | 0,26 | 0,31 | 0,25 | 0,25 | 0,29 | 0,26 | 0,27 | 0,26 | 0,34 | 0,40 | 0,07 | | | | | | | | | | | | |
| 16 | 0,31 | 0,19 | 0,24 | 0,24 | 0,26 | 0,22 | 0,24 | 0,27 | 0,25 | 0,23 | 0,26 | 0,34 | 0,44 | 0,04 | 0,14 | | | | | | | | | | | |
| 17 | 0,34 | 0,20 | 0,32 | 0,29 | 0,33 | 0,30 | 0,31 | 0,34 | 0,30 | 0,29 | 0,29 | 0,40 | 0,51 | 0,09 | 0,14 | 0,13 | | | | | | | | | | |
| 18 | 0,41 | 0,28 | 0,34 | 0,33 | 0,36 | 0,31 | 0,33 | 0,36 | 0,34 | 0,33 | 0,33 | 0,41 | 0,51 | 0,13 | 0,18 | 0,11 | 0,11 | | | | | | | | | |
| 19 | 0,39 | 0,23 | 0,30 | 0,29 | 0,33 | 0,29 | 0,31 | 0,34 | 0,30 | 0,30 | 0,30 | 0,39 | 0,50 | 0,08 | 0,12 | 0,10 | 0,09 | 0,07 | | | | | | | | |
| 20 | 0,47 | 0,31 | 0,36 | 0,38 | 0,40 | 0,34 | 0,35 | 0,38 | 0,37 | 0,35 | 0,38 | 0,46 | 0,51 | 0,17 | 0,18 | 0,16 | 0,19 | 0,16 | 0,16 | | | | | | | |
| 21 | 0,51 | 0,36 | 0,41 | 0,43 | 0,44 | 0,38 | 0,40 | 0,42 | 0,41 | 0,40 | 0,43 | 0,50 | 0,55 | 0,24 | 0,25 | 0,22 | 0,25 | 0,19 | 0,22 | 0,07 | | | | | | |
| 22 | 0,50 | 0,30 | 0,39 | 0,38 | 0,40 | 0,33 | 0,35 | 0,38 | 0,37 | 0,36 | 0,40 | 0,48 | 0,57 | 0,14 | 0,18 | 0,15 | 0,17 | 0,16 | 0,14 | 0,12 | 0,22 | | | | | |
| 23 | 0,51 | 0,32 | 0,38 | 0,39 | 0,42 | 0,33 | 0,34 | 0,37 | 0,38 | 0,36 | 0,40 | 0,48 | 0,55 | 0,10 | 0,15 | 0,10 | 0,16 | 0,11 | 0,10 | 0,07 | 0,13 | 0,07 | | | | |
| 24 | 0,41 | 0,25 | 0,32 | 0,32 | 0,35 | 0,29 | 0,30 | 0,34 | 0,32 | 0,31 | 0,33 | 0,40 | 0,47 | 0,11 | 0,16 | 0,11 | 0,13 | 0,09 | 0,10 | 0,08 | 0,10 | 0,10 | 0,06 | | | |
| 25 | 0,38 | 0,26 | 0,21 | 0,20 | 0,16 | 0,15 | 0,09 | 0,09 | 0,05 | 0,13 | 0,14 | 0,19 | 0,39 | 0,32 | 0,35 | 0,33 | 0,39 | 0,42 | 0,38 | 0,46 | 0,49 | 0,46 | 0,47 | 0,40 | | |
| 26 | 0,37 | 0,22 | 0,31 | 0,32 | 0,31 | 0,26 | 0,28 | 0,30 | 0,27 | 0,24 | 0,28 | 0,37 | 0,43 | 0,15 | 0,19 | 0,09 | 0,18 | 0,19 | 0,18 | 0,20 | 0,25 | 0,22 | 0,19 | 0,17 | 0,36 | |
| 27 | 0,57 | 0,43 | 0,48 | 0,48 | 0,48 | 0,43 | 0,46 | 0,48 | 0,48 | 0,47 | 0,50 | 0,56 | 0,65 | 0,21 | 0,32 | 0,15 | 0,26 | 0,21 | 0,20 | 0,28 | 0,31 | 0,31 | 0,22 | 0,16 | 0,54 | 0,29 |

**Supplementary table 4:** Results of GeneClass2 assignment test based on the Bayesian method of Rannala and Mountain (1997). The three first pairs of columns show, for each individual, the most likely populations and their relatives scores. Pop 9, 16 and 24 refers to natural populations of Saint-Brieuc, Atlantic Bridge, and Frøya respectively. The three following columns show -log values of the likelihoods.

| Assigned sample | Rank 1 | Score % | Rank 2 | Score % | Rank 3 | Score % | Pop 9 -Log10(L) | Pop 16 -Log10(L) | Pop 24 -Log10(L) | Nb of loci | Used loci | Missing loci |
|---|---|---|---|---|---|---|---|---|---|---|---|---|
| C-WEED_01 | Pop 9 | 100 | Pop 16 | 0 | Pop 24 | 0 | 10.728 | 29.142 | 34.966 | 21 | Sacl.11 Sacl.37 Sacl.54 Sacl.65 Sacl.95 Sacl.13 Sacl.21 Sacl.33 Sacl.41 Sacl.56 Sacl.60 Sacl.78 Sacl.88 SLN319 SLN320 SLN34 SLN35 SLN.510 SLN.32 SLN.36 SLN.54 | |
| C-WEED_02 | Pop 9 | 100 | Pop 16 | 0 | Pop 24 | 0 | 12.275 | 27.789 | 33.28 | 20 | Sacl.11 Sacl.37 Sacl.54 Sacl.95 Sacl.13 Sacl.21 Sacl.33 Sacl.41 Sacl.56 Sacl.60 Sacl.78 Sacl.88 SLN319 SLN320 SLN34 SLN35 SLN.510 SLN.32 SLN.36 SLN.54 | Sacl.65 |
| C-WEED_03 | Pop 9 | 100 | Pop 16 | 0 | Pop 24 | 0 | 7.772 | 23.752 | 33.123 | 20 | Sacl.11 Sacl.37 Sacl.54 Sacl.65 Sacl.13 Sacl.21 Sacl.33 Sacl.41 Sacl.56 Sacl.60 Sacl.78 Sacl.88 SLN319 SLN320 SLN34 SLN35 SLN.510 SLN.32 SLN.36 SLN.54 | Sacl.95 |
| C-WEED_04 | Pop 9 | 100 | Pop 16 | 0 | Pop 24 | 0 | 9.526 | 23.934 | 34.247 | 20 | Sacl.11 Sacl.37 Sacl.54 Sacl.65 Sacl.13 Sacl.21 Sacl.33 Sacl.41 Sacl.56 Sacl.60 Sacl.78 Sacl.88 SLN319 SLN320 SLN34 SLN35 SLN.510 SLN.32 SLN.36 SLN.54 | Sacl.95 |
| C-WEED_05 | Pop 9 | 100 | Pop 16 | 0 | Pop 24 | 0 | 10.625 | 18.454 | 26.344 | 21 | Sacl.11 Sacl.37 Sacl.54 Sacl.65 Sacl.95 Sacl.13 Sacl.21 Sacl.33 Sacl.41 Sacl.56 Sacl.60 Sacl.78 Sacl.88 SLN319 SLN320 SLN34 SLN35 SLN.510 SLN.32 SLN.36 SLN.54 | |
| C-WEED_06 | Pop 9 | 100 | Pop 16 | 0 | Pop 24 | 0 | 9.575 | 19.785 | 26.513 | 21 | Sacl.11 Sacl.37 Sacl.54 Sacl.65 Sacl.95 Sacl.13 Sacl.21 Sacl.33 Sacl.41 Sacl.56 Sacl.60 Sacl.78 Sacl.88 SLN319 SLN320 SLN34 SLN35 SLN.510 SLN.32 SLN.36 SLN.54 | |

| | | | | | | | | | | | | |
|---|---|---|---|---|---|---|---|---|---|---|---|---|
| C-WEED_07 | Pop 9 | 100 | Pop 16 | 0 | Pop 24 | 0 | 13.887 | 22.82 | 33.682 | 21 | Sacl.11 Sacl.37 Sacl.54 Sacl.65 Sacl.95 Sacl.13 Sacl.21 Sacl.33 Sacl.41 Sacl.56 Sacl.60 Sacl.78 Sacl.88 SLN319 SLN320 SLN34 SLN35 SLN.510 SLN.32 SLN.36 SLN.54 | |
| C-WEED_08 | Pop 9 | 100 | Pop 16 | 0 | Pop 24 | 0 | 9.882 | 24.691 | 30.887 | 21 | Sacl.11 Sacl.37 Sacl.54 Sacl.65 Sacl.95 Sacl.13 Sacl.21 Sacl.33 Sacl.41 Sacl.56 Sacl.60 Sacl.78 Sacl.88 SLN319 SLN320 SLN34 SLN35 SLN.510 SLN.32 SLN.36 SLN.54 | |
| C-WEED_09 | Pop 9 | 100 | Pop 16 | 0 | Pop 24 | 0 | 8.342 | 24.312 | 33.334 | 21 | Sacl.11 Sacl.37 Sacl.54 Sacl.65 Sacl.95 Sacl.13 Sacl.21 Sacl.33 Sacl.41 Sacl.56 Sacl.60 Sacl.78 Sacl.88 SLN319 SLN320 SLN34 SLN35 SLN.510 SLN.32 SLN.36 SLN.54 | |
| C-WEED_10 | Pop 9 | 100 | Pop 16 | 0 | Pop 24 | 0 | 10.717 | 28.346 | 35.878 | 21 | Sacl.11 Sacl.37 Sacl.54 Sacl.65 Sacl.95 Sacl.13 Sacl.21 Sacl.33 Sacl.41 Sacl.56 Sacl.60 Sacl.78 Sacl.88 SLN319 SLN320 SLN34 SLN35 SLN.510 SLN.32 SLN.36 SLN.54 | |
| C-WEED_11 | Pop 9 | 100 | Pop 16 | 0 | Pop 24 | 0 | 9.053 | 22.297 | 32.432 | 21 | Sacl.11 Sacl.37 Sacl.54 Sacl.65 Sacl.95 Sacl.13 Sacl.21 Sacl.33 Sacl.41 Sacl.56 Sacl.60 Sacl.78 Sacl.88 SLN319 SLN320 SLN34 SLN35 SLN.510 SLN.32 SLN.36 SLN.54 | |
| C-WEED_12 | Pop 9 | 100 | Pop 16 | 0 | Pop 24 | 0 | 8.656 | 22.189 | 31.243 | 18 | Sacl.11 Sacl.37 Sacl.65 Sacl.13 Sacl.21 Sacl.33 Sacl.41 Sacl.56 Sacl.60 Sacl.78 Sacl.88 SLN320 SLN34 SLN35 SLN.510 SLN.32 SLN.36 SLN.54 | Sacl.54 Sacl.95 SLN319 |
| C-WEED_13 | Pop 9 | 100 | Pop 16 | 0 | Pop 24 | 0 | 9.645 | 21.595 | 28.365 | 21 | Sacl.11 Sacl.37 Sacl.54 Sacl.65 Sacl.95 Sacl.13 Sacl.21 Sacl.33 Sacl.41 Sacl.56 Sacl.60 Sacl.78 Sacl.88 SLN319 SLN320 SLN34 SLN35 SLN.510 SLN.32 SLN.36 SLN.54 | |

| | | | | | | | | | | | |
|---|---|---|---|---|---|---|---|---|---|---|---|
| C-WEED_14 | Pop 9 | 100 | Pop 16 | 0 | Pop 24 | 0 | 9.274 | 23.635 | 31.785 | 21 | Sacl.11 Sacl.37 Sacl.54 Sacl.65 Sacl.95 Sacl.13 Sacl.21 Sacl.33 Sacl.41 Sacl.56 Sacl.60 Sacl.78 Sacl.88 SLN319 SLN320 SLN34 SLN35 SLN.510 SLN.32 SLN.36 SLN.54 |
| C-WEED_15 | Pop 9 | 100 | Pop 16 | 0 | Pop 24 | 0 | 9.496 | 24.364 | 34.102 | 21 | Sacl.11 Sacl.37 Sacl.54 Sacl.65 Sacl.95 Sacl.13 Sacl.21 Sacl.33 Sacl.41 Sacl.56 Sacl.60 Sacl.78 Sacl.88 SLN319 SLN320 SLN34 SLN35 SLN.510 SLN.32 SLN.36 SLN.54 |
| C-WEED_16 | Pop 9 | 100 | Pop 16 | 0 | Pop 24 | 0 | 10.683 | 25.698 | 33.042 | 21 | Sacl.11 Sacl.37 Sacl.54 Sacl.65 Sacl.95 Sacl.13 Sacl.21 Sacl.33 Sacl.41 Sacl.56 Sacl.60 Sacl.78 Sacl.88 SLN319 SLN320 SLN34 SLN35 SLN.510 SLN.32 SLN.36 SLN.54 |
| C-WEED_17 | Pop 9 | 100 | Pop 16 | 0 | Pop 24 | 0 | 9.093 | 19.766 | 27.755 | 21 | Sacl.11 Sacl.37 Sacl.54 Sacl.65 Sacl.95 Sacl.13 Sacl.21 Sacl.33 Sacl.41 Sacl.56 Sacl.60 Sacl.78 Sacl.88 SLN319 SLN320 SLN34 SLN35 SLN.510 SLN.32 SLN.36 SLN.54 |
| C-WEED_18 | Pop 9 | 100 | Pop 16 | 0 | Pop 24 | 0 | 11.008 | 17.141 | 24.889 | 21 | Sacl.11 Sacl.37 Sacl.54 Sacl.65 Sacl.95 Sacl.13 Sacl.21 Sacl.33 Sacl.41 Sacl.56 Sacl.60 Sacl.78 Sacl.88 SLN319 SLN320 SLN34 SLN35 SLN.510 SLN.32 SLN.36 SLN.54 |
| C-WEED_19 | Pop 9 | 100 | Pop 16 | 0 | Pop 24 | 0 | 9.977 | 24.182 | 32.448 | 21 | Sacl.11 Sacl.37 Sacl.54 Sacl.65 Sacl.95 Sacl.13 Sacl.21 Sacl.33 Sacl.41 Sacl.56 Sacl.60 Sacl.78 Sacl.88 SLN319 SLN320 SLN34 SLN35 SLN.510 SLN.32 SLN.36 SLN.54 |
| C-WEED_20 | Pop 9 | 100 | Pop 16 | 0 | Pop 24 | 0 | 10.24 | 26.503 | 33.171 | 21 | Sacl.11 Sacl.37 Sacl.54 Sacl.65 Sacl.95 Sacl.13 Sacl.21 Sacl.33 Sacl.41 Sacl.56 Sacl.60 Sacl.78 Sacl.88 SLN319 SLN320 SLN34 SLN35 SLN.510 SLN.32 SLN.36 SLN.54 |

| | | | | | | | | | | | | |
|---|---|---|---|---|---|---|---|---|---|---|---|---|
| C-WEED_21 | Pop 9 | 100 | Pop 16 | 0 | Pop 24 | 0 | 8.868 | 23.543 | 31.715 | 21 | Sacl.11 Sacl.37 Sacl.54 Sacl.65 Sacl.95 Sacl.13 Sacl.21 Sacl.33 Sacl.41 Sacl.56 Sacl.60 Sacl.78 Sacl.88 SLN319 SLN320 SLN34 SLN35 SLN.510 SLN.32 SLN.36 SLN.54 | |
| C-WEED_22 | Pop 9 | 100 | Pop 16 | 0 | Pop 24 | 0 | 9.789 | 23.174 | 29.135 | 20 | Sacl.11 Sacl.37 Sacl.65 Sacl.95 Sacl.13 Sacl.21 Sacl.33 Sacl.41 Sacl.56 Sacl.60 Sacl.78 Sacl.88 SLN319 SLN320 SLN34 SLN35 SLN.510 SLN.32 SLN.36 SLN.54 | Sacl.54 |
| C-WEED_23 | Pop 9 | 100 | Pop 16 | 0 | Pop 24 | 0 | 9.743 | 22.02 | 28.364 | 20 | Sacl.11 Sacl.37 Sacl.54 Sacl.65 Sacl.95 Sacl.13 Sacl.21 Sacl.33 Sacl.56 Sacl.60 Sacl.78 Sacl.88 SLN319 SLN320 SLN34 SLN35 SLN.510 SLN.32 SLN.36 SLN.54 | Sacl.41 |
| C-WEED_24 | Pop 9 | 100 | Pop 16 | 0 | Pop 24 | 0 | 10.512 | 29.472 | 36.816 | 21 | Sacl.11 Sacl.37 Sacl.54 Sacl.65 Sacl.95 Sacl.13 Sacl.21 Sacl.33 Sacl.41 Sacl.56 Sacl.60 Sacl.78 Sacl.88 SLN319 SLN320 SLN34 SLN35 SLN.510 SLN.32 SLN.36 SLN.54 | |
| C-WEED_25 | Pop 9 | 100 | Pop 16 | 0 | Pop 24 | 0 | 8.592 | 19.767 | 27.957 | 21 | Sacl.11 Sacl.37 Sacl.54 Sacl.65 Sacl.95 Sacl.13 Sacl.21 Sacl.33 Sacl.41 Sacl.56 Sacl.60 Sacl.78 Sacl.88 SLN319 SLN320 SLN34 SLN35 SLN.510 SLN.32 SLN.36 SLN.54 | |
| C-WEED_26 | Pop 9 | 100 | Pop 16 | 0 | Pop 24 | 0 | 9.641 | 24.874 | 32.439 | 21 | Sacl.11 Sacl.37 Sacl.54 Sacl.65 Sacl.95 Sacl.13 Sacl.21 Sacl.33 Sacl.41 Sacl.56 Sacl.60 Sacl.78 Sacl.88 SLN319 SLN320 SLN34 SLN35 SLN.510 SLN.32 SLN.36 SLN.54 | |
| C-WEED_27 | Pop 9 | 100 | Pop 16 | 0 | Pop 24 | 0 | 10.587 | 28.777 | 36.219 | 21 | Sacl.11 Sacl.37 Sacl.54 Sacl.65 Sacl.95 Sacl.13 Sacl.21 Sacl.33 Sacl.41 Sacl.56 Sacl.60 Sacl.78 Sacl.88 SLN319 SLN320 SLN34 SLN35 SLN.510 SLN.32 SLN.36 SLN.54 | |

| | | | | | | | | | | | | |
|---|---|---|---|---|---|---|---|---|---|---|---|---|
| C-WEED_28 | Pop 9 | 100 | Pop 16 | 0 | Pop 24 | 0 | 9.992 | 22.785 | 32.291 | 21 | Sacl.11 Sacl.37 Sacl.54 Sacl.65 Sacl.95 Sacl.13 Sacl.21 Sacl.33 Sacl.41 Sacl.56 Sacl.60 Sacl.78 Sacl.88 SLN319 SLN320 SLN34 SLN35 SLN.510 SLN.32 SLN.36 SLN.54 | |
| C-WEED_29 | Pop 9 | 100 | Pop 16 | 0 | Pop 24 | 0 | 9.021 | 21.81 | 31.362 | 21 | Sacl.11 Sacl.37 Sacl.54 Sacl.65 Sacl.95 Sacl.13 Sacl.21 Sacl.33 Sacl.41 Sacl.56 Sacl.60 Sacl.78 Sacl.88 SLN319 SLN320 SLN34 SLN35 SLN.510 SLN.32 SLN.36 SLN.54 | |
| C-WEED_30 | Pop 9 | 100 | Pop 16 | 0 | Pop 24 | 0 | 10.897 | 22.605 | 27.718 | 21 | Sacl.11 Sacl.37 Sacl.54 Sacl.65 Sacl.95 Sacl.13 Sacl.21 Sacl.33 Sacl.41 Sacl.56 Sacl.60 Sacl.78 Sacl.88 SLN319 SLN320 SLN34 SLN35 SLN.510 SLN.32 SLN.36 SLN.54 | |
| C-WEED_31 | Pop 9 | 100 | Pop 16 | 0 | Pop 24 | 0 | 10.092 | 26.912 | 34.372 | 21 | Sacl.11 Sacl.37 Sacl.54 Sacl.65 Sacl.95 Sacl.13 Sacl.21 Sacl.33 Sacl.41 Sacl.56 Sacl.60 Sacl.78 Sacl.88 SLN319 SLN320 SLN34 SLN35 SLN.510 SLN.32 SLN.36 SLN.54 | |
| C-WEED_32 | Pop 9 | 100 | Pop 16 | 0 | Pop 24 | 0 | 9.275 | 20.628 | 24.17 | 17 | Sacl.11 Sacl.37 Sacl.95 Sacl.13 Sacl.21 Sacl.33 Sacl.56 Sacl.60 Sacl.88 SLN319 SLN320 SLN34 SLN35 SLN.510 SLN.32 SLN.36 SLN.54 | Sacl.54 Sacl.65 Sacl.41 Sacl.78 |
| C-WEED_33 | Pop 9 | 100 | Pop 16 | 0 | Pop 24 | 0 | 8.11 | 23.222 | 31.466 | 21 | Sacl.11 Sacl.37 Sacl.54 Sacl.65 Sacl.95 Sacl.13 Sacl.21 Sacl.33 Sacl.41 Sacl.56 Sacl.60 Sacl.78 Sacl.88 SLN319 SLN320 SLN34 SLN35 SLN.510 SLN.32 SLN.36 SLN.54 | |
| C-WEED_34 | Pop 9 | 100 | Pop 16 | 0 | Pop 24 | 0 | 9.113 | 21.116 | 26.905 | 21 | Sacl.11 Sacl.37 Sacl.54 Sacl.65 Sacl.95 Sacl.13 Sacl.21 Sacl.33 Sacl.41 Sacl.56 Sacl.60 Sacl.78 Sacl.88 SLN319 SLN320 SLN34 SLN35 SLN.510 SLN.32 SLN.36 SLN.54 | |
| C-WEED_35 | Pop 9 | 100 | Pop 16 | 0 | Pop 24 | 0 | 12.398 | 26.945 | 32.887 | 21 | Sacl.11 Sacl.37 Sacl.54 Sacl.65 Sacl.95 Sacl.13 Sacl.21 Sacl.33 | |

| | | | | | | | | | | | | |
|---|---|---|---|---|---|---|---|---|---|---|---|---|
| | | | | | | | | | | | Sacl.41 Sacl.56 Sacl.60 Sacl.78 Sacl.88 SLN319 SLN320 SLN34 SLN35 SLN.510 SLN.32 SLN.36 SLN.54 | |
| SAMS_10 | Pop 16 | 100 | Pop 9 | 0 | Pop 24 | 0 | 26.206 | 18.044 | 31.694 | 21 | Sacl.11 Sacl.37 Sacl.54 Sacl.65 Sacl.95 Sacl.13 Sacl.21 Sacl.33 Sacl.41 Sacl.56 Sacl.60 Sacl.78 Sacl.88 SLN319 SLN320 SLN34 SLN35 SLN.510 SLN.32 SLN.36 SLN.54 | |
| SAMS_11 | Pop 16 | 100 | Pop 24 | 0 | Pop 9 | 0 | 30.636 | 20.466 | 28.669 | 21 | Sacl.11 Sacl.37 Sacl.54 Sacl.65 Sacl.95 Sacl.13 Sacl.21 Sacl.33 Sacl.41 Sacl.56 Sacl.60 Sacl.78 Sacl.88 SLN319 SLN320 SLN34 SLN35 SLN.510 SLN.32 SLN.36 SLN.54 | |
| SAMS_12 | Pop 16 | 100 | Pop 24 | 0 | Pop 9 | 0 | 24.579 | 15.724 | 23.815 | 21 | Sacl.11 Sacl.37 Sacl.54 Sacl.65 Sacl.95 Sacl.13 Sacl.21 Sacl.33 Sacl.41 Sacl.56 Sacl.60 Sacl.78 Sacl.88 SLN319 SLN320 SLN34 SLN35 SLN.510 SLN.32 SLN.36 SLN.54 | |
| SAMS_13 | Pop 16 | 100 | Pop 24 | 0 | Pop 9 | 0 | 22.534 | 13.551 | 21.84 | 18 | Sacl.11 Sacl.37 Sacl.54 Sacl.95 Sacl.13 Sacl.21 Sacl.33 Sacl.56 Sacl.60 Sacl.78 SLN319 SLN320 SLN34 SLN35 SLN.510 SLN.32 SLN.36 SLN.54 | Sacl.65 Sacl.41 Sacl.88 |
| SAMS_14 | Pop 16 | 100 | Pop 9 | 0 | Pop 24 | 0 | 27.426 | 19.877 | 28.444 | 21 | Sacl.11 Sacl.37 Sacl.54 Sacl.65 Sacl.95 Sacl.13 Sacl.21 Sacl.33 Sacl.41 Sacl.56 Sacl.60 Sacl.78 Sacl.88 SLN319 SLN320 SLN34 SLN35 SLN.510 SLN.32 SLN.36 SLN.54 | |
| SAMS_16 | Pop 16 | 99.989 | Pop 24 | 0.011 | Pop 9 | 0 | 27.909 | 17.557 | 21.532 | 21 | Sacl.11 Sacl.37 Sacl.54 Sacl.65 Sacl.95 Sacl.13 Sacl.21 Sacl.33 Sacl.41 Sacl.56 Sacl.60 Sacl.78 Sacl.88 SLN319 SLN320 SLN34 SLN35 SLN.510 SLN.32 SLN.36 SLN.54 | |
| SAMS_17 | Pop 16 | 99.998 | Pop 24 | 0.001 | Pop 9 | 0.001 | 19.877 | 14.821 | 19.849 | 19 | Sacl.37 Sacl.54 Sacl.95 Sacl.13 Sacl.21 Sacl.33 Sacl.41 Sacl.56 Sacl.60 Sacl.78 Sacl.88 SLN319 | Sacl.11 Sacl.65 |

| | | | | | | | | | | | | |
|---|---|---|---|---|---|---|---|---|---|---|---|---|
| | | | | | | | | | | | SLN320 SLN34 SLN35 SLN.510 SLN.32 SLN.36 SLN.54 | |
| SAMS_18 | Pop 16 | 99.999 | Pop 9 | 0.001 | Pop 24 | 0 | 22.558 | 17.72 | 26.629 | 21 | Sacl.11 Sacl.37 Sacl.54 Sacl.65 Sacl.95 Sacl.13 Sacl.21 Sacl.33 Sacl.41 Sacl.56 Sacl.60 Sacl.78 Sacl.88 SLN319 SLN320 SLN34 SLN35 SLN.510 SLN.32 SLN.36 SLN.54 | |
| SAMS_19 | Pop 16 | 100 | Pop 24 | 0 | Pop 9 | 0 | 30.053 | 16.046 | 25.332 | 20 | Sacl.11 Sacl.37 Sacl.54 Sacl.65 Sacl.95 Sacl.13 Sacl.21 Sacl.33 Sacl.41 Sacl.56 Sacl.78 Sacl.88 SLN319 SLN320 SLN34 SLN35 SLN.510 SLN.32 SLN.36 SLN.54 | Sacl.60 |
| SAMS_21 | Pop 16 | 100 | Pop 9 | 0 | Pop 24 | 0 | 28.371 | 15.833 | 29.609 | 21 | Sacl.11 Sacl.37 Sacl.54 Sacl.65 Sacl.95 Sacl.13 Sacl.21 Sacl.33 Sacl.41 Sacl.56 Sacl.60 Sacl.78 Sacl.88 SLN319 SLN320 SLN34 SLN35 SLN.510 SLN.32 SLN.36 SLN.54 | |
| SAMS_22 | Pop 16 | 100 | Pop 24 | 0 | Pop 9 | 0 | 34.009 | 14.09 | 21.748 | 18 | Sacl.11 Sacl.37 Sacl.54 Sacl.65 Sacl.95 Sacl.13 Sacl.33 Sacl.41 Sacl.56 Sacl.88 SLN319 SLN320 SLN34 SLN35 SLN.510 SLN.32 SLN.36 SLN.54 | Sacl.21 Sacl.60 Sacl.78 |
| SAMS_23 | Pop 16 | 99.998 | Pop 9 | 0.002 | Pop 24 | 0 | 22.329 | 17.633 | 28.138 | 21 | Sacl.11 Sacl.37 Sacl.54 Sacl.65 Sacl.95 Sacl.13 Sacl.21 Sacl.33 Sacl.41 Sacl.56 Sacl.60 Sacl.78 Sacl.88 SLN319 SLN320 SLN34 SLN35 SLN.510 SLN.32 SLN.36 SLN.54 | |
| SAMS_24 | Pop 16 | 100 | Pop 24 | 0 | Pop 9 | 0 | 30.214 | 17.863 | 24.81 | 20 | Sacl.11 Sacl.37 Sacl.54 Sacl.65 Sacl.95 Sacl.13 Sacl.21 Sacl.33 Sacl.56 Sacl.60 Sacl.78 Sacl.88 SLN319 SLN320 SLN34 SLN35 SLN.510 SLN.32 SLN.36 SLN.54 | Sacl.41 |
| SAMS_25 | Pop 16 | 100 | Pop 24 | 0 | Pop 9 | 0 | 25.206 | 14.096 | 22.2 | 19 | Sacl.11 Sacl.37 Sacl.65 Sacl.95 Sacl.13 Sacl.21 Sacl.33 Sacl.41 Sacl.56 Sacl.60 Sacl.78 Sacl.88 | Sacl.54 SLN34 |

| | | | | | | | | | | | | |
|---|---|---|---|---|---|---|---|---|---|---|---|---|
| | | | | | | | | | | | SLN319 SLN320 SLN35 SLN.510 SLN.32 SLN.36 SLN.54 | |
| SAMS_26 | Pop 16 | 100 | Pop 24 | 0 | Pop 9 | 0 | 26.988 | 17.079 | 22.864 | 20 | Sacl.11 Sacl.37 Sacl.54 Sacl.65 Sacl.95 Sacl.13 Sacl.21 Sacl.33 Sacl.41 Sacl.56 Sacl.78 Sacl.88 SLN319 SLN320 SLN34 SLN35 SLN.510 SLN.32 SLN.36 SLN.54 | Sacl.60 |
| SAMS_27 | Pop 16 | 99.799 | Pop 24 | 0.201 | Pop 9 | 0 | 20.236 | 14.117 | 16.814 | 15 | Sacl.11 Sacl.37 Sacl.95 Sacl.21 Sacl.33 Sacl.41 Sacl.56 Sacl.78 SLN319 SLN320 SLN34 SLN.510 SLN.32 SLN.36 SLN.54 | Sacl.54 Sacl.65 Sacl.13 Sacl.60 Sacl.88 SLN35 |
| SAMS_28 | Pop 16 | 100 | Pop 24 | 0 | Pop 9 | 0 | 28.132 | 13.718 | 20.254 | 21 | Sacl.11 Sacl.37 Sacl.54 Sacl.65 Sacl.95 Sacl.13 Sacl.21 Sacl.33 Sacl.41 Sacl.56 Sacl.60 Sacl.78 Sacl.88 SLN319 SLN320 SLN34 SLN35 SLN.510 SLN.32 SLN.36 SLN.54 | |
| SAMS_29 | Pop 16 | 100 | Pop 9 | 0 | Pop 24 | 0 | 22.401 | 15.056 | 28.312 | 21 | Sacl.11 Sacl.37 Sacl.54 Sacl.65 Sacl.95 Sacl.13 Sacl.21 Sacl.33 Sacl.41 Sacl.56 Sacl.60 Sacl.78 Sacl.88 SLN319 SLN320 SLN34 SLN35 SLN.510 SLN.32 SLN.36 SLN.54 | |
| SAMS_30 | Pop 16 | 100 | Pop 9 | 0 | Pop 24 | 0 | 21.178 | 14.663 | 23.523 | 16 | Sacl.11 Sacl.37 Sacl.54 Sacl.13 Sacl.21 Sacl.33 Sacl.56 Sacl.60 SLN319 SLN320 SLN34 SLN35 SLN.510 SLN.32 SLN.36 SLN.54 | Sacl.65 Sacl.95 Sacl.41 Sacl.78 Sacl.88 |
| SAMS_31 | Pop 16 | 99.989 | Pop 24 | 0.011 | Pop 9 | 0 | 29.895 | 16.232 | 20.209 | 18 | Sacl.11 Sacl.37 Sacl.54 Sacl.65 Sacl.95 Sacl.13 Sacl.21 Sacl.33 Sacl.41 Sacl.56 Sacl.60 Sacl.88 SLN320 SLN34 SLN.510 SLN.32 SLN.36 SLN.54 | Sacl.78 SLN319 SLN35 |
| SAMS_32 | Pop 16 | 99.999 | Pop 9 | 0.001 | Pop 24 | 0 | 24.63 | 19.66 | 31.865 | 21 | Sacl.11 Sacl.37 Sacl.54 Sacl.65 Sacl.95 Sacl.13 Sacl.21 Sacl.33 Sacl.41 Sacl.56 Sacl.60 Sacl.78 | |

| | | | | | | | | | | | | |
|---|---|---|---|---|---|---|---|---|---|---|---|---|
| | | | | | | | | | | | Sacl.88 SLN319 SLN320 SLN34 SLN35 SLN.510 SLN.32 SLN.36 SLN.54 | |
| SAMS_33 | Pop 16 | 100 | Pop 9 | 0 | Pop 24 | 0 | 17.752 | 11.78 | 22.727 | 18 | Sacl.11 Sacl.37 Sacl.95 Sacl.13 Sacl.21 Sacl.33 Sacl.41 Sacl.56 Sacl.60 Sacl.78 SLN319 SLN320 SLN34 SLN35 SLN.510 SLN.32 SLN.36 SLN.54 | Sacl.54 Sacl.65 Sacl.88 |
| SAMS_34 | Pop 16 | 100 | Pop 24 | 0 | Pop 9 | 0 | 22.663 | 13.685 | 20.438 | 19 | Sacl.37 Sacl.54 Sacl.95 Sacl.13 Sacl.21 Sacl.33 Sacl.41 Sacl.56 Sacl.60 Sacl.78 Sacl.88 SLN319 SLN320 SLN34 SLN35 SLN.510 SLN.32 SLN.36 SLN.54 | Sacl.11 Sacl.65 |
| SAMS_35 | Pop 16 | 100 | Pop 9 | 0 | Pop 24 | 0 | 22.857 | 13.292 | 26.938 | 21 | Sacl.11 Sacl.37 Sacl.54 Sacl.65 Sacl.95 Sacl.13 Sacl.21 Sacl.33 Sacl.41 Sacl.56 Sacl.60 Sacl.78 Sacl.88 SLN319 SLN320 SLN34 SLN35 SLN.510 SLN.32 SLN.36 SLN.54 | |
| SAMS_36 | Pop 16 | 100 | Pop 24 | 0 | Pop 9 | 0 | 28.657 | 19.043 | 28.021 | 20 | Sacl.11 Sacl.37 Sacl.54 Sacl.65 Sacl.13 Sacl.21 Sacl.33 Sacl.41 Sacl.56 Sacl.60 Sacl.78 Sacl.88 SLN319 SLN320 SLN34 SLN35 SLN.510 SLN.32 SLN.36 SLN.54 | Sacl.95 |
| SES_01 | Pop 16 | 91.825 | Pop 24 | 8.175 | Pop 9 | 0 | 34.464 | 15.001 | 16.052 | 21 | Sacl.11 Sacl.37 Sacl.54 Sacl.65 Sacl.95 Sacl.13 Sacl.21 Sacl.33 Sacl.41 Sacl.56 Sacl.60 Sacl.78 Sacl.88 SLN319 SLN320 SLN34 SLN35 SLN.510 SLN.32 SLN.36 SLN.54 | |
| SES_02 | Pop 24 | 60.029 | Pop 16 | 39.971 | Pop 9 | 0 | 37.693 | 16.044 | 15.868 | 21 | Sacl.11 Sacl.37 Sacl.54 Sacl.65 Sacl.95 Sacl.13 Sacl.21 Sacl.33 Sacl.41 Sacl.56 Sacl.60 Sacl.78 Sacl.88 SLN319 SLN320 SLN34 SLN35 SLN.510 SLN.32 SLN.36 SLN.54 | |
| SES_03 | Pop 24 | 99.579 | Pop 16 | 0.421 | Pop 9 | 0 | 38.56 | 15.3 | 12.926 | 21 | Sacl.11 Sacl.37 Sacl.54 Sacl.65 Sacl.95 Sacl.13 Sacl.21 Sacl.33 Sacl.41 Sacl.56 Sacl.60 Sacl.78 | |

| | | | | | | | | | | | |
|---|---|---|---|---|---|---|---|---|---|---|---|
| | | | | | | | | | | | Sacl.88 SLN319 SLN320 SLN34 SLN35 SLN.510 SLN.32 SLN.36 SLN.54 |
| SES_04 | Pop 16 | 59.763 | Pop 24 | 40.237 | Pop 9 | 0 | 30.571 | 12.592 | 12.764 | 21 | Sacl.11 Sacl.37 Sacl.54 Sacl.65 Sacl.95 Sacl.13 Sacl.21 Sacl.33 Sacl.41 Sacl.56 Sacl.60 Sacl.78 Sacl.88 SLN319 SLN320 SLN34 SLN35 SLN.510 SLN.32 SLN.36 SLN.54 |
| SES_05 | Pop 24 | 79.129 | Pop 16 | 20.871 | Pop 9 | 0 | 31.648 | 11.957 | 11.378 | 21 | Sacl.11 Sacl.37 Sacl.54 Sacl.65 Sacl.95 Sacl.13 Sacl.21 Sacl.33 Sacl.41 Sacl.56 Sacl.60 Sacl.78 Sacl.88 SLN319 SLN320 SLN34 SLN35 SLN.510 SLN.32 SLN.36 SLN.54 |
| SES_06 | Pop 24 | 97.479 | Pop 16 | 2.521 | Pop 9 | 0 | 33.238 | 13.322 | 11.735 | 21 | Sacl.11 Sacl.37 Sacl.54 Sacl.65 Sacl.95 Sacl.13 Sacl.21 Sacl.33 Sacl.41 Sacl.56 Sacl.60 Sacl.78 Sacl.88 SLN319 SLN320 SLN34 SLN35 SLN.510 SLN.32 SLN.36 SLN.54 |
| SES_07 | Pop 24 | 99.872 | Pop 16 | 0.128 | Pop 9 | 0 | 30.4 | 13.616 | 10.724 | 21 | Sacl.11 Sacl.37 Sacl.54 Sacl.65 Sacl.95 Sacl.13 Sacl.21 Sacl.33 Sacl.41 Sacl.56 Sacl.60 Sacl.78 Sacl.88 SLN319 SLN320 SLN34 SLN35 SLN.510 SLN.32 SLN.36 SLN.54 |
| SES_08 | Pop 24 | 99.999 | Pop 16 | 0.001 | Pop 9 | 0 | 33.544 | 15.494 | 10.538 | 21 | Sacl.11 Sacl.37 Sacl.54 Sacl.65 Sacl.95 Sacl.13 Sacl.21 Sacl.33 Sacl.41 Sacl.56 Sacl.60 Sacl.78 Sacl.88 SLN319 SLN320 SLN34 SLN35 SLN.510 SLN.32 SLN.36 SLN.54 |
| SES_09 | Pop 24 | 93.122 | Pop 16 | 6.878 | Pop 9 | 0 | 30.461 | 12.42 | 11.289 | 21 | Sacl.11 Sacl.37 Sacl.54 Sacl.65 Sacl.95 Sacl.13 Sacl.21 Sacl.33 Sacl.41 Sacl.56 Sacl.60 Sacl.78 Sacl.88 SLN319 SLN320 SLN34 SLN35 SLN.510 SLN.32 SLN.36 SLN.54 |
| SES_10 | Pop 24 | 99.908 | Pop 16 | 0.092 | Pop 9 | 0 | 39.714 | 18.971 | 15.935 | 21 | Sacl.11 Sacl.37 Sacl.54 Sacl.65 Sacl.95 Sacl.13 Sacl.21 Sacl.33 Sacl.41 Sacl.56 Sacl.60 Sacl.78 |

| | | | | | | | | | | | | |
|---|---|---|---|---|---|---|---|---|---|---|---|---|
| | | | | | | | | | | | Sacl.88 SLN319 SLN320 SLN34 SLN35 SLN.510 SLN.32 SLN.36 SLN.54 | |
| SES_11 | Pop 16 | 96.933 | Pop 24 | 3.067 | Pop 9 | 0 | 33.867 | 11.585 | 13.085 | 21 | Sacl.11 Sacl.37 Sacl.54 Sacl.65 Sacl.95 Sacl.13 Sacl.21 Sacl.33 Sacl.41 Sacl.56 Sacl.60 Sacl.78 Sacl.88 SLN319 SLN320 SLN34 SLN35 SLN.510 SLN.32 SLN.36 SLN.54 | |
| SES_12 | Pop 16 | 91.825 | Pop 24 | 8.175 | Pop 9 | 0 | 34.464 | 15.001 | 16.052 | 21 | Sacl.11 Sacl.37 Sacl.54 Sacl.65 Sacl.95 Sacl.13 Sacl.21 Sacl.33 Sacl.41 Sacl.56 Sacl.60 Sacl.78 Sacl.88 SLN319 SLN320 SLN34 SLN35 SLN.510 SLN.32 SLN.36 SLN.54 | |
| SES_13 | Pop 24 | 99.582 | Pop 16 | 0.418 | Pop 9 | 0 | 34.062 | 13.429 | 11.052 | 21 | Sacl.11 Sacl.37 Sacl.54 Sacl.65 Sacl.95 Sacl.13 Sacl.21 Sacl.33 Sacl.41 Sacl.56 Sacl.60 Sacl.78 Sacl.88 SLN319 SLN320 SLN34 SLN35 SLN.510 SLN.32 SLN.36 SLN.54 | |
| SES_14 | Pop 16 | 100 | Pop 24 | 0 | Pop 9 | 0 | 28.906 | 11.437 | 17.218 | 21 | Sacl.11 Sacl.37 Sacl.54 Sacl.65 Sacl.95 Sacl.13 Sacl.21 Sacl.33 Sacl.41 Sacl.56 Sacl.60 Sacl.78 Sacl.88 SLN319 SLN320 SLN34 SLN35 SLN.510 SLN.32 SLN.36 SLN.54 | |
| SES_15 | Pop 16 | 85.177 | Pop 24 | 14.823 | Pop 9 | 0 | 30.247 | 12.395 | 13.154 | 21 | Sacl.11 Sacl.37 Sacl.54 Sacl.65 Sacl.95 Sacl.13 Sacl.21 Sacl.33 Sacl.41 Sacl.56 Sacl.60 Sacl.78 Sacl.88 SLN319 SLN320 SLN34 SLN35 SLN.510 SLN.32 SLN.36 SLN.54 | |
| SES_16 | Pop 24 | 99.438 | Pop 16 | 0.562 | Pop 9 | 0 | 32.728 | 15.233 | 12.985 | 21 | Sacl.11 Sacl.37 Sacl.54 Sacl.65 Sacl.95 Sacl.13 Sacl.21 Sacl.33 Sacl.41 Sacl.56 Sacl.60 Sacl.78 Sacl.88 SLN319 SLN320 SLN34 SLN35 SLN.510 SLN.32 SLN.36 SLN.54 | |
| SES_17 | Pop 24 | 96.081 | Pop 16 | 3.919 | Pop 9 | 0 | 29.873 | 14.009 | 12.619 | 20 | Sacl.37 Sacl.54 Sacl.65 Sacl.95 Sacl.13 Sacl.21 Sacl.33 Sacl.41 Sacl.56 Sacl.60 Sacl.78 Sacl.88 | Sacl.11 |

| | | | | | | | | | | | |
|---|---|---|---|---|---|---|---|---|---|---|---|
| | | | | | | | | | | | SLN319 SLN320 SLN34 SLN35 SLN.510 SLN.32 SLN.36 SLN.54 |
| SES_18 | Pop 24 | 99.944 | Pop 16 | 0.056 | Pop 9 | 0 | 34.079 | 14.476 | 11.223 | 21 | Sacl.11 Sacl.37 Sacl.54 Sacl.65 Sacl.95 Sacl.13 Sacl.21 Sacl.33 Sacl.41 Sacl.56 Sacl.60 Sacl.78 Sacl.88 SLN319 SLN320 SLN34 SLN35 SLN.510 SLN.32 SLN.36 SLN.54 |
| SES_19 | Pop 24 | 99.651 | Pop 16 | 0.349 | Pop 9 | 0 | 33.527 | 13.261 | 10.805 | 21 | Sacl.11 Sacl.37 Sacl.54 Sacl.65 Sacl.95 Sacl.13 Sacl.21 Sacl.33 Sacl.41 Sacl.56 Sacl.60 Sacl.78 Sacl.88 SLN319 SLN320 SLN34 SLN35 SLN.510 SLN.32 SLN.36 SLN.54 |
| SES_20 | Pop 16 | 86.78 | Pop 24 | 13.22 | Pop 9 | 0 | 30.237 | 14.048 | 14.865 | 21 | Sacl.11 Sacl.37 Sacl.54 Sacl.65 Sacl.95 Sacl.13 Sacl.21 Sacl.33 Sacl.41 Sacl.56 Sacl.60 Sacl.78 Sacl.88 SLN319 SLN320 SLN34 SLN35 SLN.510 SLN.32 SLN.36 SLN.54 |
| SES_21 | Pop 24 | 93.411 | Pop 16 | 6.589 | Pop 9 | 0 | 37.574 | 14.582 | 13.431 | 21 | Sacl.11 Sacl.37 Sacl.54 Sacl.65 Sacl.95 Sacl.13 Sacl.21 Sacl.33 Sacl.41 Sacl.56 Sacl.60 Sacl.78 Sacl.88 SLN319 SLN320 SLN34 SLN35 SLN.510 SLN.32 SLN.36 SLN.54 |
| SES_22 | Pop 24 | 99.993 | Pop 16 | 0.007 | Pop 9 | 0 | 31.865 | 14.092 | 9.908 | 21 | Sacl.11 Sacl.37 Sacl.54 Sacl.65 Sacl.95 Sacl.13 Sacl.21 Sacl.33 Sacl.41 Sacl.56 Sacl.60 Sacl.78 Sacl.88 SLN319 SLN320 SLN34 SLN35 SLN.510 SLN.32 SLN.36 SLN.54 |
| SES_23 | Pop 16 | 89.144 | Pop 24 | 10.856 | Pop 9 | 0 | 33.439 | 11.531 | 12.446 | 21 | Sacl.11 Sacl.37 Sacl.54 Sacl.65 Sacl.95 Sacl.13 Sacl.21 Sacl.33 Sacl.41 Sacl.56 Sacl.60 Sacl.78 Sacl.88 SLN319 SLN320 SLN34 SLN35 SLN.510 SLN.32 SLN.36 SLN.54 |
| SES_24 | Pop 24 | 99.991 | Pop 16 | 0.009 | Pop 9 | 0 | 29.847 | 15.76 | 11.691 | 21 | Sacl.11 Sacl.37 Sacl.54 Sacl.65 Sacl.95 Sacl.13 Sacl.21 Sacl.33 Sacl.41 Sacl.56 Sacl.60 Sacl.78 |

| | | | | | | | | | | | |
|---|---|---|---|---|---|---|---|---|---|---|---|
| | | | | | | | | | | | Sacl.88 SLN319 SLN320 SLN34 SLN35 SLN.510 SLN.32 SLN.36 SLN.54 |
| SES_25 | Pop 16 | 91.524 | Pop 24 | 8.476 | Pop 9 | 0 | 24.789 | 11.959 | 12.992 | 21 | Sacl.11 Sacl.37 Sacl.54 Sacl.65 Sacl.95 Sacl.13 Sacl.21 Sacl.33 Sacl.41 Sacl.56 Sacl.60 Sacl.78 Sacl.88 SLN319 SLN320 SLN34 SLN35 SLN.510 SLN.32 SLN.36 SLN.54 |
| SES_26 | Pop 16 | 98.903 | Pop 24 | 1.097 | Pop 9 | 0 | 33.868 | 12.296 | 14.251 | 21 | Sacl.11 Sacl.37 Sacl.54 Sacl.65 Sacl.95 Sacl.13 Sacl.21 Sacl.33 Sacl.41 Sacl.56 Sacl.60 Sacl.78 Sacl.88 SLN319 SLN320 SLN34 SLN35 SLN.510 SLN.32 SLN.36 SLN.54 |
| SES_27 | Pop 24 | 99.761 | Pop 16 | 0.239 | Pop 9 | 0 | 30.686 | 15.843 | 13.223 | 21 | Sacl.11 Sacl.37 Sacl.54 Sacl.65 Sacl.95 Sacl.13 Sacl.21 Sacl.33 Sacl.41 Sacl.56 Sacl.60 Sacl.78 Sacl.88 SLN319 SLN320 SLN34 SLN35 SLN.510 SLN.32 SLN.36 SLN.54 |
| SES_28 | Pop 24 | 99.995 | Pop 16 | 0.005 | Pop 9 | 0 | 30.171 | 15.312 | 11.029 | 21 | Sacl.11 Sacl.37 Sacl.54 Sacl.65 Sacl.95 Sacl.13 Sacl.21 Sacl.33 Sacl.41 Sacl.56 Sacl.60 Sacl.78 Sacl.88 SLN319 SLN320 SLN34 SLN35 SLN.510 SLN.32 SLN.36 SLN.54 |
| SES_29 | Pop 16 | 83.677 | Pop 24 | 16.323 | Pop 9 | 0 | 31.038 | 11.778 | 12.488 | 21 | Sacl.11 Sacl.37 Sacl.54 Sacl.65 Sacl.95 Sacl.13 Sacl.21 Sacl.33 Sacl.41 Sacl.56 Sacl.60 Sacl.78 Sacl.88 SLN319 SLN320 SLN34 SLN35 SLN.510 SLN.32 SLN.36 SLN.54 |
| SES_30 | Pop 24 | 93.634 | Pop 16 | 6.366 | Pop 9 | 0 | 36.423 | 13.195 | 12.027 | 21 | Sacl.11 Sacl.37 Sacl.54 Sacl.65 Sacl.95 Sacl.13 Sacl.21 Sacl.33 Sacl.41 Sacl.56 Sacl.60 Sacl.78 Sacl.88 SLN319 SLN320 SLN34 SLN35 SLN.510 SLN.32 SLN.36 SLN.54 |
